# Post-Ictal Gamma Oscillations Predict Hippocampal Structural Integrity in Mesial Temporal Lobe Epilepsy

**DOI:** 10.1101/2025.02.02.636164

**Authors:** Daniel J. Valdivia, Fabio Cesar Tescarollo, Koray Ercan, Isaac Huang, Betsy Vasquez, Lauren Parsonnet, Sangmi Chung, Detlev Boison, Robert Gross, Ezequiel Gleichgerrcht, Spencer C. Chen, Hai Sun

## Abstract

The underlying histopathology and epileptic network of patients with mesial temporal lobe epilepsy (mTLE) are difficult to ascertain. Here, we report a novel electrical activity following seizure termination, termed post-ictal gamma oscillations (PIGOs), recorded in mice with intrahippocampal kainic acid injection and a patient with mTLE. PIGOs are characterized by a spectral shift of increased power in gamma frequencies relative to decreased power in other frequencies, generating a gamma peak. PIGOs are accompanied by increased intracellular calcium levels among parvalbumin-positive interneurons and a direct current shift. Only a subgroup of animals in the study had PIGOs. These animals had less pronounced hippocampal sclerosis (HS) than those without PIGOs. To illustrate the translational potential of these findings, we analyzed data from two patients with unilateral mTLE, one with PIGOs and the other without. The patient with PIGOs had symmetrical hippocampi on neuroimaging, whereas the other exhibited overly decreased interictal glucose uptake in left hippocampus, suggesting left-sided HS. After receiving laser ablation of mesial temporal regions, the patient with PIGOs became seizure-free, whereas the other did not. Our results suggest that PIGOs may serve as a biomarker for a milder form of HS in patients with mTLE and for predicting treatment outcomes.

## Introduction

Epilepsy is a widespread debilitating condition affecting approximately 1% of the world’s population, with mesial temporal lobe epilepsy (mTLE) being the most common type of focal epilepsy among adults (1–5). Among patients with mTLE, hippocampal sclerosis (HS), characterized by atrophy and hardening of the hippocampal tissue, is the most frequent histopathologic abnormality encountered in the seizure onset zone (SOZ) (6–8). These abnormalities can be attributed to pyramidal cell loss, granular cell dispersion, and astrogliosis in various subregions of the Ammon’s horn (6–8). HS can often be detected with neuroimaging studies such as magnetic resonance imaging (MRI) or 18-fluoro-deoxyglucose positron emission tomography (FDG-PET) (9–12). When a patient with drug-resistant mTLE is being considered for surgical intervention, the finding of imaging suggestive of HS can predict better seizure control with surgical resection or laser ablation of the sclerotic hippocampus (11, 13, 14). However, a portion of mTLE patients may not show classic hippocampal abnormalities on preoperative imaging evaluation. In these cases, the implantation of intracranial electrodes may be necessary to identify the SOZ (15). Since localizing the SOZ can be challenging even with the intracranial recordings, it is desirable to identify biomarkers capable of predicting underlying tissue pathology and guiding treatment decisions for patients with mTLE.

Since seizures commonly self-terminate, most epilepsy research has focused on seizure initiation (16–22), with less emphasis on the electrophysiological and cellular mechanisms involved in seizure termination and the post-ictal state. Seizure termination may occur through various mechanisms, including local processes such as local adenosine triphosphate and glutamate depletion, glial buffering, increased gamma-aminobutyric acid (GABA) activity, as well as remote mechanisms like the increased activity of thalamic and brainstem nuclei (3, 23–25). Moreover, seizure termination is commonly followed by a phenomenon known as the post-ictal state, characterized by prolonged post-ictal generalized electroencephalographic suppression (PGES). PGES has been linked to an increased risk of cardiorespiratory arrest and sudden unexpected death in epilepsy (SUDEP) (26–30). Focal epilepsy, such as mTLE, has been increasingly viewed as a brain network pathology (31–33). A better understanding of seizure termination and post-ictal state will also help delineate the epileptic network, develop novel therapies to shorten seizures, and reduce morbidity and mortality associated with epilepsy.

In this study, we identified and characterized a stereotyped pattern of gamma-range oscillatory electrical activity following the termination of spontaneous seizures, termed post-ictal gamma oscillations (PIGOs), recorded from a murine model of mTLE resulting from the intrahippocampal kainic acid (IHKA) injection. PIGOs had distinct spectral characteristics from the preceding seizure. Most importantly, PIGOs were only observed in the postictal period among a subset of animals included in this study, referred to as PIGO(+) animals. Subsequent evaluation of hippocampal histology revealed significant differences in hippocampal architecture between PIGO(+) and PIGO(−) animals. Finally, we showed two patients with medically intractable unilateral mTLE, one with and one without PIGOs following seizures, and compared the neuroimaging findings of these two patients, highlighting the clinical relevance of our findings from the murine model of mTLE.

## Results

### Post-ictal gamma oscillations emerge in a subset of epileptic animals

The intrahippocampal kainic acid (IHKA) murine model, which produces electrophysiological and histopathological features of human mTLE (34–36) , was used in this study. The 16 animals included in this study were divided into two experimental groups with regard to the anteroposterior and dorsoventral position where the IHKA injections were made in the hippocampus: (i) animals that had kainic acid (KA) injected in the anterodorsal hippocampal CA1 layer (adIHKA, n=8; Figure 1Ai), and (ii) animals in which KA was injected in the CA1 layer of the posteroventral portion of the hippocampus (pvIHKA, n=8; Figure 1Aii). This distinction was made to have the EEG recorded from variable positions in the dorsal hippocampus relative to the IHKA injection site: ipsilateral to the IHKA injection (iCA1), contralateral to the IHKA injection (cCA1), dentate gyrus ipsilateral to IHKA injection (iDG), and contralateral to IHKA injection (cDG). Detailed descriptions of surgical techniques can be found in Methods.

**Figure 1.**
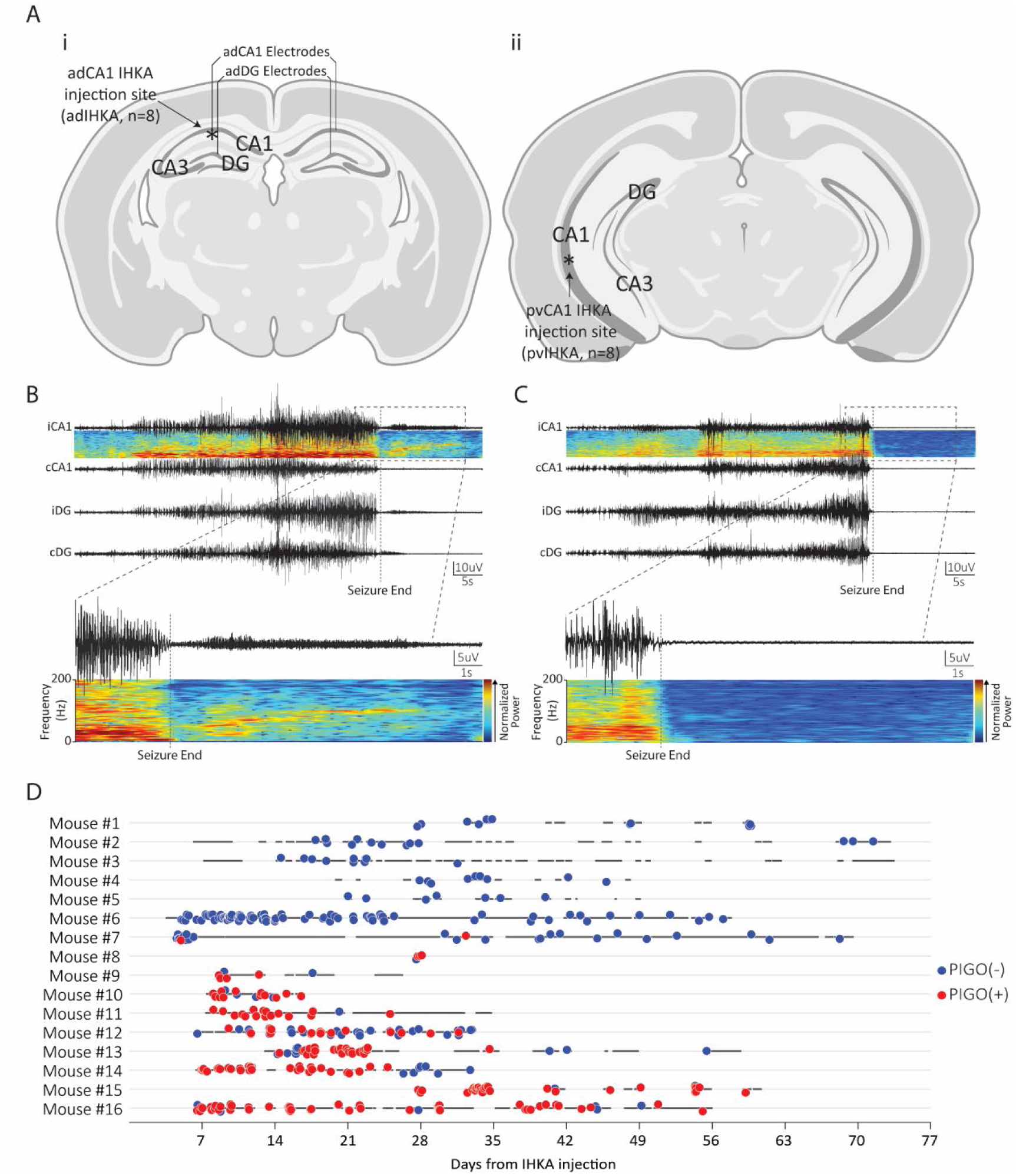
PIGOs were identified in a subset of epileptic animals. **A**, A schematic diagram showing, in **i**, the locations of the KA injection site in adIHKA-injected mice, and recording electrodes implantation sites, and in **ii**, the KA injection site in pvIHKA-injected mice. **B**, Sample EEG recording of a spontaneous seizure PIGOs identified in the dorsal iCA1, iDG and cDG recording channels within the hippocampus. Note the increased gamma-range activity following seizure termination in the spectrogram for iCA1. **C**, As in **B**, but for a PIGO(-)Sz. Note the complete EEG suppression following seizure termination observed in the spectrogram in the iCA1. **D**, Timeline of experiment relative to time (days) after initial KA injection of all mice included in this study (n=16). PIGO(+)Sz (red circles), PIGO(-)Sz (blue circles) and days where animals were recorded are indicated (Black dots) are also represented.

Mice were recorded up to 75 days after the IHKA injection. To assess the presence of PIGOs, only spontaneously occurring generalized seizures with motor components were included in the analysis, resulting in 416 seizures. Among the seizures detected, PIGOs were identified based on two specific criteria: 1) narrow-band activity superimposed on the post-ictal depression in the spectrogram (later characterized in this study as gamma-range activity), and 2) a correlation between the narrow-band activity and a crescendo-decrescendo pattern in the amplitude of oscillatory activity in the LFP (Figure 1B). For comparison, a seizure without PIGOs is shown in Figure 1C.

Among 416 seizures recorded from 16 animals, 189 (45.4%) seizures were followed by PIGOs and deemed PIGO-positive seizures (PIGO(+)Sz), while the remaining 227 (54.6%) seizures were not followed by PIGOs and deemed PIGO-negative seizures (PIGO(-)Sz). Ten (62.5%) animals in this study had one or more PIGO(+)Szs, while six (37.5%) displayed only PIGO(-)Szs. They were denoted as PIGO(+) and PIGO(-) animals, respectively. Among PIGO(+) animals, the mean incidence of PIGO(+)Sz was 69% (ranging from 8 to 94%). The incidence of PIGO(+)Szs and PIGO(-)Szs remained unchanged over the entire recording period (Figure 1D). A similar seizure frequency was observed between groups (Supplementary Figures 1A and 1C), but PIGO(+) animals exhibited a lower average seizure frequency than PIGO(-) animals after weeks 3-4 of the seizure timeline (Supplementary Figure 1B).

### The pvIHKA animals exhibited more PIGOs than adIHKA animals

A key finding of this study is that PIGO(+)Szs were only recorded in a subset of animals (Figure 1D). When grouped by IHKA injection site, more animals with pvIHKA exhibited PIGOs than animals with adIHKA (7/8 - 87.5% and 3/8 - 37.5%, respectively; *p*=0.12; Figure 2A). pvIHKA animals also showed a significantly higher cumulative percentage of PIGO(+)Szs compared to adIHKA animals, both when all animals were included (adIHKA, n=8; pvIHKA, n=8; *p*<0.01), and when only PIGO(+) animals were analyzed (adIHKA, n=3; pvIHKA, n=7; *p*<0.05; Figures 2B and 2C, respectively). Additionally, the incidence of PIGOs was greater among the pvIHKA group compared to the adIHKA group (*p*=0.08; Figure 2D). Since the objective of this study is to characterize the electrophysiological properties of PIGO and its underlying cellular and network mechanisms, we performed the subsequent analyses by comparing PIGO(+) with PIGO(-) animals.

**Figure 2.**
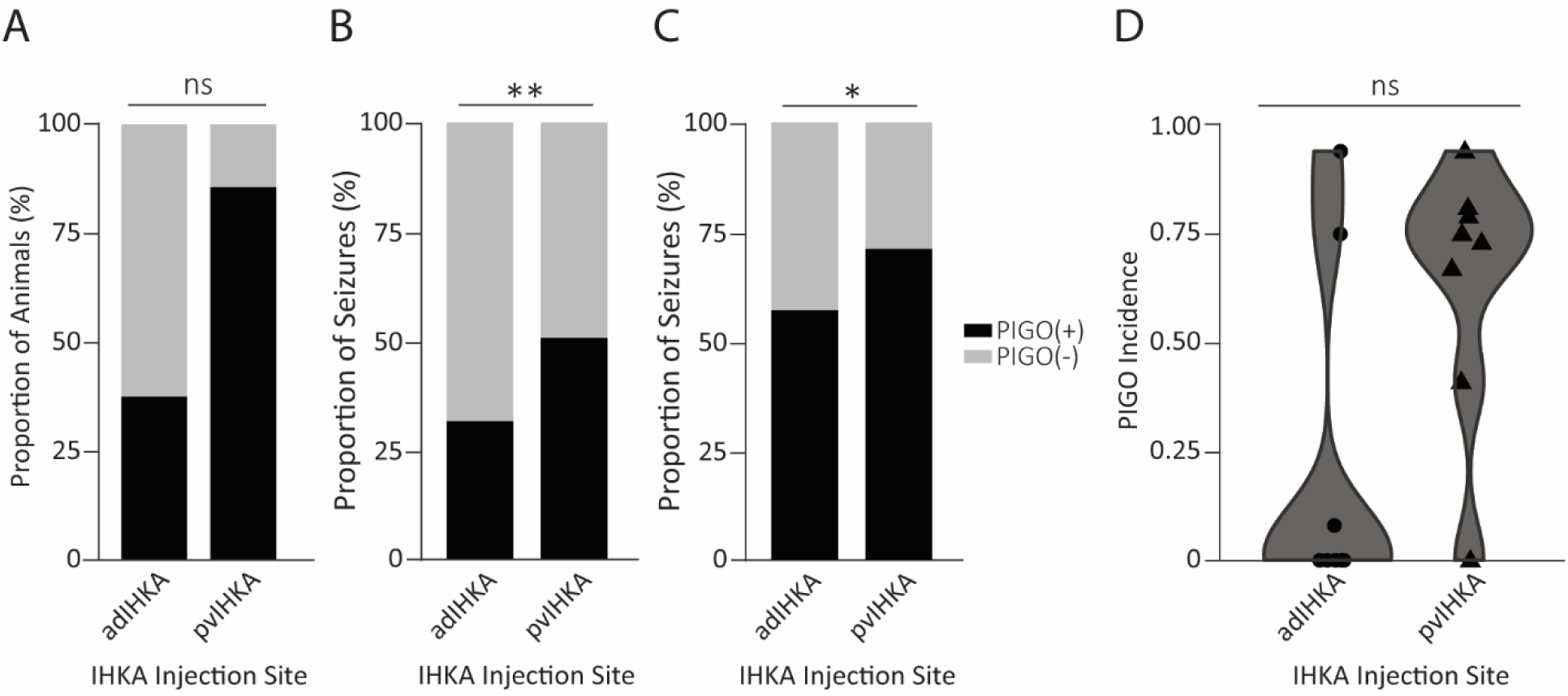
Relative position of IHKA injection influences PIGOs incidence. **A,** Proportion of animals injected with IHKA in either the antero-dorsal (adIHKA) or posterior-ventral (pvIHKA) CA1 layer that presented PIGOs following seizures (3/8 animals from the adIHKA group - 37.5%; 7/8 animals from the pvIHKA group – 87.5%; P=0.12; Fisher’s exact test). **B**, Cumulative proportion of PIGO(+)Sz in relation to the total number of seizures observed in each experimental group (adIHKA group, n=8: 36/114 = 32.5%; pvIHKA group, n=8: 153/302 = 51.0%, **P<0.01; Pearson’s Chi-squared test). **C**, Proportion of PIGO(+)Sz in relation to the total number of seizures observed in PIGO(+) animals (adIHKA group, n=3: 36/63 = 57.1%; pvIHKA group, n=7: 153/215 = 71.2%, **P<0.05; Pearson’s Chi-squared test). **D**, PIGO incidence (number of PIGO(+)Sz / total number of seizures) for each animal from the adIHKA (n=8) and pvIHKA groups (n=8). The adIHKA and pvIHKA group had an incidence of 0.22 ± 0.14 (range: 0 – 0.94) and 0.64 ± 0.11 (range: 0 – 0.94), respectively (P=0.08; Wilcoxon rank sum test). Note that while 75.00% of pvIHKA animals (n=6/8) displayed PIGOs in more than 50% of seizures, 62.5% of the adIHKA-injected animals (n=5/8) did not display any PIGOs. Within the adIHKA group, one animal displayed PIGOs in 8.00% of seizures (n=2/26), whereas the remaining two animals displayed PIGOs in more than 50% of the observed seizures.

### Behavioral and electrophysiological characteristics of PIGOs

To assess and compare the behavioral characteristics and manifestations associated with PIGO(+) and PIGO(-)Szs, we implemented a modified Racine Score (RS), adapted from *Erum et al*. (37), and assigned behavioral scores to all seizures in the study (n=416). When all animals were included in the analysis, no significant difference in seizure severity was observed between PIGO(+)Szs and PIGO(-)Szs (*p*=0.7, Supplementary Figure 1D). However, among PIGO(+) animals, the average RS of PIGO(+)Szs was lower than PIGO(-)Szs (*p*<0.01, Supplementary Figure 1E). PIGO(+)Szs were also, on average, shorter than PIGO(-)Szs (*p*<0.05; Supplementary Figure 1F). We also analyzed and assigned an additional behavioral score on the post-ictal EEG of PIGO(+)Sz. During the postictal period where PIGOs were found, animal behavior was consistent with the postictal state, exhibiting motor arrest, occasionally followed by facial automatisms. The average RS during PIGOs was 1.43 (median: 1, range: 0–4).

Sparingly, PIGOs were more likely to appear on the side of the hippocampus that received IHKA injection (*p*=0.07; Figure 3Ai and 3Aii). Furthermore, PIGO(+)Szs rarely displayed PIGOs only in the contralateral hemisphere, as the majority of seizures showed PIGOs bilaterally or ipsilaterally (Figure 3B). Principal component analysis (PCA) was used to unwrap the distribution of PIGOs among the four recorded hippocampal sublayers (Table 1, Figure 3C). PCA analysis revealed three clusters of recording channel dominance: the first cluster (yellow dots) was characterized by a dominance of the iCA1 (iCA1, iCA1 + cCA1, iCA1 + cCA1 + iDG); the second cluster (purple dots) by the dominance of iDG (iDG, iDG + iCA1, iDG + iCA1 + cDG); the third grouping (teal dots) by PIGOs occurring in all four recording channels.

**Figure 3.**
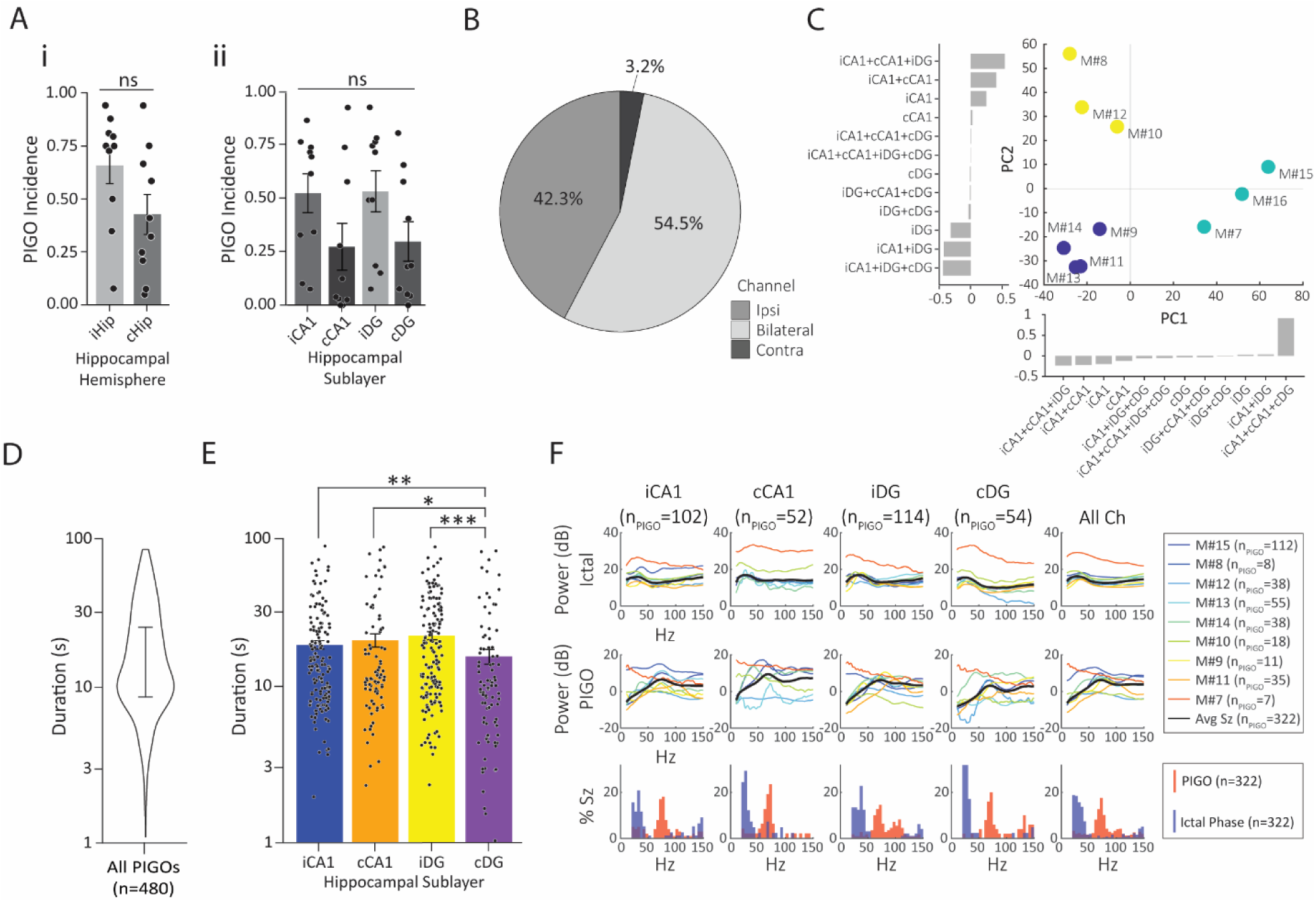
PIGOs location, duration, power, and frequency. **A**, Incidence rates of PIGO(+)Sz for PIGO(+) animals (n=10) segregated by (i) hemisphere and (ii) individual hippocampal sublayers. Note that the incidence rate is higher in the ipsilateral hemisphere (iHip = 0.65 ± 0.09, cHip = 0.43 ± 0.09; P=0.07; Wilcoxon rank-sum test) and in hippocampal sublayers ipsilateral to the IHKA injection (iCA1 = 0.53 ± 0.09, cCA1 = 0.28 ± 0.11, iDG = 0.54 ± 0.09, cDG = 0.30 ± 0.09; iCA1 vs. cCA1: p>0.05; iDG vs. cDG: P>0.05, One-Way ANOVA with Tukey’s Multiple comparisons test as *post hoc*). **B**, Proportion of seizures (n=189) from PIGO(+) animals exhibiting PIGOs exclusively on ipsilateral or contralateral recording electrodes or any combination of ipsilateral and contralateral channels (bilateral). **C**, Principal component analysis with K-means clustering of the PIGO channel combinations for each animal (n=10). **D**, Duration of all PIGOs (n=480) grouped (mean(± SEM)): 18.30s ± 0.68 ; median: 12.44s; range: 1.02s-75.18s; IQR: 8.3s-23.5s). **E**, PIGO durations subdivided by hippocampal sublayer (mean ± (SEM)): iCA1 = 17.74s ± 1.08 (range: 1.93s - 75.18s; median: 13.31s; IQR: 1.93s – 75.2s), cCA1 = 18.97s ± 1.81 (range: 2.30s - 74.22s; median: 11.32s; IQR: 9.04s - 22.73s), iDG = 20.30s ± 1.18 (range: 2.30s - 73.74s; median: 14.97s; IQR: 8.74s - 29.38s), and cDG = 14.97s ± 1.60 (range: 5.75s - 16.85s; median: 9.64s; IQR: 5.75s – 16.85s) (iCA1 vs. cDG: **P<0.01; cCA1 vs. cDG: *P<0.05; iDG vs. cDG: ***P<0.0005; Two-Way ANOVA with Tukey’s Multiple comparisons test as *post hoc*). ANOVA for durations computed in logarithmic space. **F**, Power spectrum of the ictal phase (top row) and PIGO phase (middle row) with peak frequencies plotted (bottom row). In total, 322 PIGO(+) EEG traces from 142 seizures were included. The upper row refers to the power spectrum (in Db) of the ictal period in the different frequencies (in Hz) in the different recording channels (iCA1, cCA1, iDG, cDG) and in all channels altogether (last column). Middle Row, same as the upper row, but for the PIGO phase of seizures. The bottom row shows the peak frequency occupancy of PIGOs (red bars) or the seizures ictal phase (blue bars). Peak frequencies were significantly different between the ictal phase and PIGO phase across all channels (iCA1: P<0.0001; cCA1: P<0.0001; iDG: P<0.0001; cDG: P<0.0001), as well as cumulatively (All channels: P<0.0001; Wilcoxon rank-sum test).

**Table 1.** Proportion of seizures with PIGOs delineated by unique channel combination of PIGO emergence.

| Mouse ID | Seizures (n) | PIGO(+) Sz (n) | iCA1 | iDG | cCA1 | cDG | iCA1+iDG | iCA1+cCA1 | iCA1+cDG | iDG+cDG | iDG+cCA1 | cCA1+cDG | iCA1/iDG+cDG | iCA1+cCA1+iDG | iCA1+cCA1+cDG | iDG+cCA1+cDG | iCA1+cCA1+iDG+cDG |
| --- | --- | --- | --- | --- | --- | --- | --- | --- | --- | --- | --- | --- | --- | --- | --- | --- | --- |
| Mouse 15 | 33 | 31 | - | - | 6.45 | - | - | 6.45 | - | - | - | - | - | - | 6.45 | - | 80.65 |
| Mouse 8 | 4 | 3 | - | - | - | - | - | 33.33 | - | - | - | - | - | 66.67 | - | - | - |
| Mouse 1 | 12 | 0 | - | - | - | - | - | - | - | - | - | - | - | - | - | - | - |
| Mouse 2 | 13 | 0 | - | - | - | - | - | - | - | - | - | - | - | - | - | - | - |
| Mouse 3 | 10 | 0 | - | - | - | - | - | - | - | - | - | - | - | - | - | - | - |
| Mouse 4 | 9 | 0 | - | - | - | - | - | - | - | - | - | - | - | - | - | - | - |
| Mouse 5 | 7 | 0 | - | - | - | - | - | - | - | - | - | - | - | - | - | - | - |
| Mouse 12 | 46 | 19 | 10.53 | - | 5.26 | 10.53 | 10.53 | 36.84 | - | - | - | - | - | 26.32 | - | - | - |
| Mouse 13 | 33 | 24 | - | - | - | - | 70.83 | - | - | - | - | - | 25.00 | 4.17 | - | - | - |
| Mouse 16 | 58 | 47 | 4.26 | 4.26 | - | - | 19.15 | - | - | - | - | - | - | - | - | - | 72.34 |
| Mouse 14 | 39 | 31 | - | 80.65 | - | - | 12.90 | - | - | 3.23 | - | - | - | - | - | 3.23 | - |
| Mouse 10 | 16 | 12 | 66.67 | - | - | - | - | 8.33 | - | 16.67 | - | - | - | - | - | - | 8.33 |
| Mouse 9 | 6 | 4 | - | - | - | 25.00 | - | - | - | 25.00 | - | - | 50.00 | - | - | - | - |
| Mouse 11 | 17 | 16 | - | 18.75 | - | - | 37.50 | - | - | 6.25 | - | - | 37.50 | - | - | - | - |
| Mouse 6 | 87 | 0 | - | - | - | - | - | - | - | - | - | - | - | - | - | - | - |
| Mouse 7 | 26 | 2 | - | - | - | - | - | - | - | - | - | - | 50.00 | - | - | - | 50.00 |

We also analyzed PIGO duration. Among the 189 PIGO(+)Szs, 480 PIGOs were identified across the four recording channels. The average PIGO duration was 18.3s (± 0.68; Figure 3D), and PIGOs identified from the cDG were the shortest (*p*<0.05; Figure 3E).

Next, we characterized the spectral characteristics of PIGOs (see Methods) and found a distinct spectral pattern compared to the preceding ictal period. For this, we analyzed 322 PIGO events across all four recording channels. For this analysis, one animal exhibited anomalous activity in two recording channels which affected the frequency characterization of the EEG. Excluding only those two channels misrepresented the averages, so the mouse was excluded altogether. These signals, however, did not interfere with the other analysis to characterize PIGO. The pre-ictal baselines were subtracted to compare ictal and post-ictal phases across seizures. The average ictal spectrogram showed relatively flat and evenly distributed power, with a slight peak below 30Hz. However, the average PIGO spectrogram showed a spectral shift into higher frequencies, with a relative decrease in power below 30Hz but a concentration of power in the gamma range with an average peak around 60-70Hz. Sorting each seizure individually, the majority of the PIGO segment’s peak frequencies fell within the gamma range (30-80Hz), constituting 56.83% of the peaks, in contrast to only 23.81% of peaks during the ictal segment. By contrast, only 9.63% of the PIGO segment’s peak frequencies were observed below 30 Hz, while 59.93% of peak frequencies in the ictal segment were below 30 Hz (Figure 3F).

### PIGOs were anticorrelated

We next investigated the correlation of PIGO activity across all four recording channels. A sample of a PIGO(+)Sz with PIGOs identified in all four recording channels is shown (Figures 4Ai and 4Aii). To quantitatively characterize this phenomenon, we computed Pearson’s correlation coefficient among all channel pairs during the post-ictal periods for all seizures recorded during our experiments. 340 seizures were included: 215 PIGO(-)Szs and 125 PIGO(+)Szs, which resulted in a total of 2040 channel pairs. Correlations were first grouped based on the presence of PIGOs in neither channel (Zero PIGO, 1,462 pairs), one channel (One PIGO, 334 pairs), or two channels (Two PIGO, 244 pairs). We observed that channel pairs with Two PIGOs showed the most anticorrelation, while channel pairs with Zero PIGOs showed slightly positive correlation (*p*<0.01; Figure 4B). When correlations were further grouped based on various channel pair combinations, the iCA1 and cCA1 channel pair exhibited the most anticorrelation (*p*<0.01; Figure 4C).

**Figure 4.**
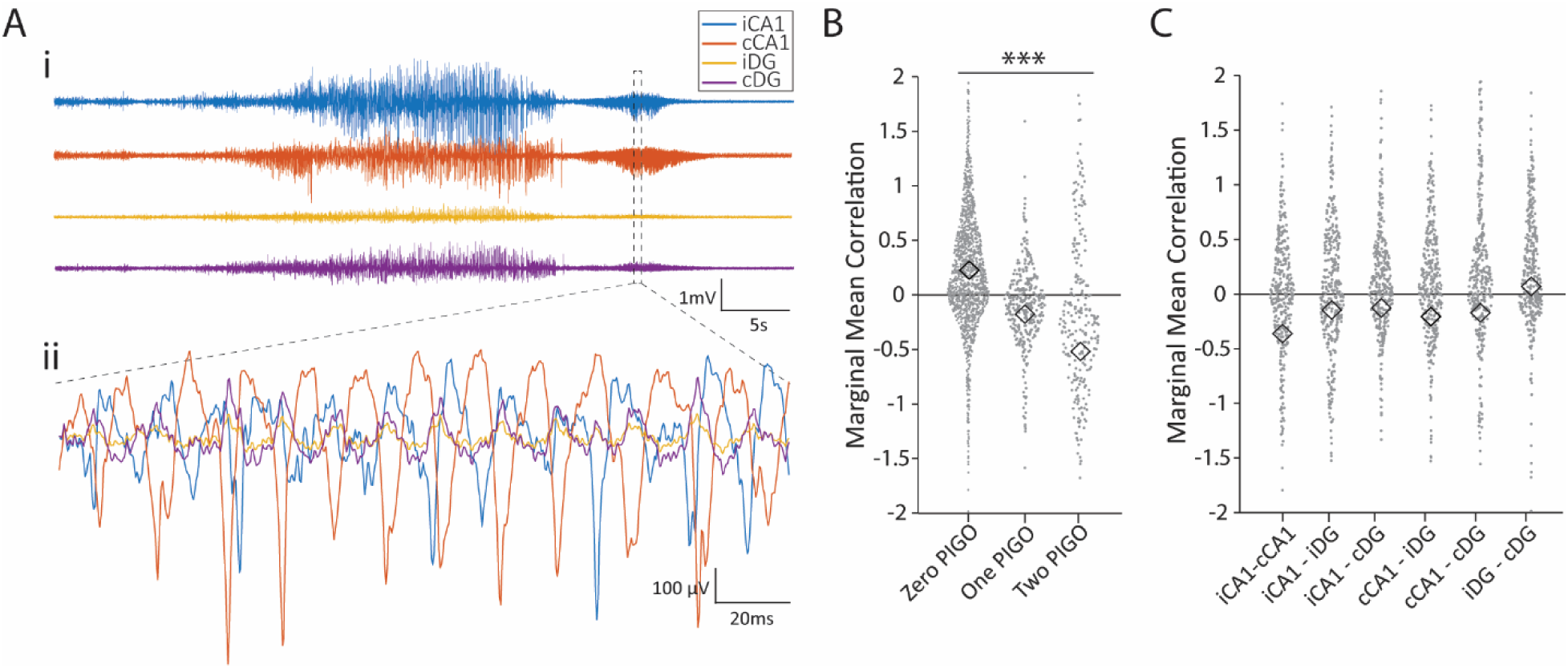
PIGO synchronicity across recording channels. **A**, **(i)** Visualization of a seizure with PIGOs present following seizure termination in all four recording channels (iCA1, cCA1, iDG, cDG), (**ii)** PIGOs from all four recording channels overlayed to show the variability between channels in terms of periodicity, amplitude, and sinusoidal activity. **B**, Synchronicity analysis of channel pairs during the PIGO time segment demonstrating the raw data distribution and marginal means of Fisher Z-Transformed and baseline-subtracted Pearson’s correlation coefficient. The standard errors were very small and omitted for visual clarity. The correlation coefficient was significantly different based on the number of PIGOs present in a correlation. Marginal means of correlation by PIGO Presence (mean(SE)): Zero PIGO (n=1462): -0.52 ± 0.13, One PIGO (n=334): -0.18 ± 0.08, Two PIGO (n=244) 0.23 ± 0.02. ***P<0.001 (Four-way ANOVA). (Zero PIGO vs. One PIGO: ***p<0.001; One PIGO vs. Two PIGO: ***p<0.001; Zero PIGO vs. Two PIGO: ***p<0.001; Four-Way ANOVA with Tukey’s multiple comparisons test as *post hoc.* The horizontal line marked with three asterisks indicates that all pairwise comparisons were statistically significant. **C**, as in **B**, with correlations (n=2040) grouped based on channel combinations. The strongest negative correlation was observed between bilateral iCA1 and cCA1 (r = -0.36 ± 0.05). The iCA1-cCA1 channel pair was significantly more negative than all other channel pairs, except for cCA1-iDG (p=0.055). The iDG-cDG channel pairing was significantly more positive than all other channel pairs. Marginal means of correlation by channel pair (mean(SE)): iCA1-cCA1: -0.36 ± 0.05, iCA1-iDG: -0.14 ± 0.05, iCA1-cDG: -0.12 ± 0.05, cCA1-iDG: -0.20 ± 0.05, cCA1-cDG: -0.17 ± 0.06, iDG-cDG: 0.07 ± 0.05. ***P<0.001 (Four-way ANOVA). (iCA1-cCA1 vs. iCA1-iDG: ***p<0.001; iCA1-cCA1 vs. iCA1-cDG: ***p<0.001; iCA1-cCA1 vs. cCA1-iDG: p>0.05; iCA1-cCA1 vs cCA1-cDG: *p<0.05; iCA1-cCA1 vs. iDG-cDG: ***p<0.001; iCA1-iDG vs. iCA1-cDG: p>0.05; iCA1-iDG vs. cCA1-iDG: p>0.05; iCA1-iDG vs. cCA1-cDG: p>0.05; iCA1-iDG vs. iDG-cDG: ***p<0.001; iCA1-cDG vs. cCA1-iDG: p>0.05; iCA1-cDG vs cCA1-cDG: p>0.05; iCA1-cDG vs. iDG-cDG: **p<0.01; cCA1-iDG vs. cCA1-cDG: p>0.05; cCA1-iDG vs iDG-cDG: ***p<0.001; cCA1-cDG vs. IDG-cDG: **p<0.01; Four-Way ANOVA with Tukey’s multiple comparisons test as *post hoc*).

### PIGOs were associated with increased intracellular Ca^2+^ of parvalbumin-positive interneurons (PV-INs)

Next, we investigated the cellular activities associated with the emergence of PIGOs using *in vivo* Ca^2+^ fiber photometry in the bilateral dorsal hippocampi of pvIHKA animals (PV-cre animals, n=6). This strategy aimed to simultaneously detect the intracellular Ca^2+^ activities ([Ca^2+^]_i_) from pyramidal neurons and PV-INs from bilateral CA1 regions (Figure 5A; Supplementary Figures 2A and 2B). Among the six animals recorded for Ca^2+^ fiber photometry, five had PIGO(+)Szs and were included in the analysis. The four [Ca^2+^]_i_ signals recorded among pyramidal neurons and PV-INs were denoted as iCaMKIIα (CaMKIIα-positive pyramidal neurons ipsilateral to the IHKA injection), iPV (PV-INs ipsilateral to the IHKA injection), cCaMKIIα (CaMKIIα-positive pyramidal neurons contralateral to the IHKA injection), and cPV (PV-INs in the CA1 area contralateral to the IHKA injection). Deflections in photometry signals (ΔF/F) were averaged across PIGO(+)Szs (n=30) or PIGO(-)Szs (n=10). The EEG activity levels acquired in the recording areas were quantified by calculating the coastline index (CI) (38, 39).

**Figure 5.**
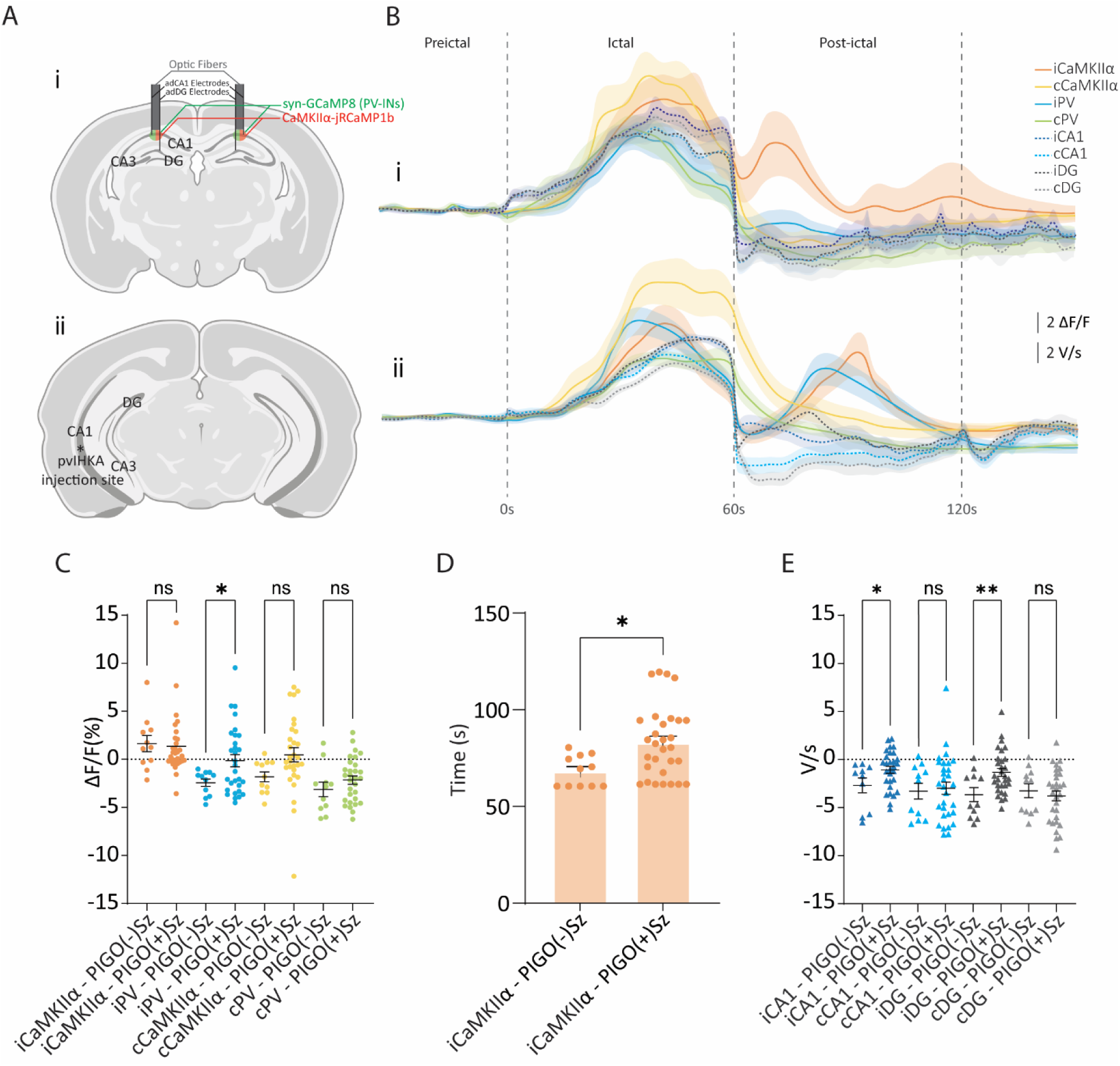
Ca^2+^ fiber photometry activity and coastline index analysis. **A**, A schematic diagram showing, in **i**, the locations of the Ca^2+^ fiber photometry optic fibers, recording electrodes implantation sites, the adeno-associated viral vectors CaMKIIα-RCaMP and syn-GCaMP (green-and-red circles), and in **ii**, the KA injection site in vIHKA-injected mice. **B**, Averaged Ca^2+^ fiber photometry (solid lines) and CI (dashed lines) activity profiles of PIGO(-)Sz, in **i**, and PIGO(+)Sz, in **ii**. **C,** Analysis of Ca^2+^ activity levels (ΔF/F) comparing each photometry signal (iCaMKIIα, iPV, cCaMKIIα, and cPV) obtained in the post-ictal period (iCaMKIIα-PIGO(-)Sz = 1.00 ± 0.61, iCaMKIIα-PIGO(+)Sz = 1.35 ± 0.58, P=0.74; iPV-PIGO(-)Sz = -2.46 ± 0.42, iPV-PIGO(+)Sz = - 0.14 ± 0.64, P<0.05; cCaMKIIα-PIGO(-)Sz = -1.97 ± 0.54, cCaMKIIα-PIGO(+)Sz = 0.47 ± 0.73, P=0.07; cPV-PIGO(-)Sz = -3.21 ± 0.83, cPV-PIGO(+)Sz = -2.16 ± 0.42, P=0.23; Unpaired t test). **D,** Timing difference of the peaks detected in the iCaMKIIα Ca^2+^ fiber photometry signals in the post-ictal period among PIGO(+)Szs (n=30) or PIGO(-)Szs (n=10) (PIGO(+)Sz: t= 68.3 ± 8.0 s, PIGO(-)Sz: t= 83.0 ± 3.3 s, P<0.05; Unpaired t-test). **E**, Analysis of CI levels comparing the averaged CI from each EEG signal (iCA1, cCA1, iDG, and cDG) recorded in the first 30 seconds of the post-ictal period of PIGO(-)Sz and PIGO(+)Sz (iCA1-PIGO(-)Sz = -2.67 ± 0.77 V/s, iCA1-PIGO(+)Sz = -1.07 ± 0.36 V/s, P<0.05; cCA1-PIGO(-)Sz = -3.52 ± 0.88 V/s, cCA1-PIGO(+)Sz = -3.00 ± 0.62 V/s, P=0.66; iDG-PIGO(-)Sz = -3.82 ± 0.78 V/s, iDG-PIGO(+)Sz = -1.34 ± 0.41 V/s, P<0.01; cDG-PIGO(-)Sz = -3.45 ± 0.76 V/s, cDG-PIGO(+)Sz = -3.78 ± 0.51, P=0.74; Unpaired t-test).

During the ictal period, the [Ca^2+^]_i_ activities among pyramidal neurons and PV-INs in bilateral CA1 displayed similar profiles in both PIGO(-)Szs and PIGO(+)Szs (*p*>0.99; Figure 5B, Supplementary Figure 2C). Similar results were observed in the EEG activities estimated using CI in bilateral CA1 and DG (iCaMKIIα: *p*=0.97, iPV: *p*=0.99, cCaMKIIα: *p*=0.99, and cPV: *p=*0.57; Figure 5B, Supplementary Figure 2D).

Similarly, prominent rises in the iCaMKIIα levels were observed during the post-ictal period in both PIGO(+)Szs and PIGO(-)Szs (Figure 5B). Although the levels of activity of iCaMKIIα did not differ between PIGO(+) and PIGO(-)Sz (*p*=0.98; Figure 5C), iCaMKIIα activities reached the peak following PIGO(-)Szs significantly earlier than among PIGO(+)Szs (*p*<0.01; Figure 5D). In addition to the rise in [Ca^2+^]_i_-iCaMKIIα levels during the post-ictal periods, in PIGO(+)Szs, we observed increased [Ca^2+^]_i_ activity in iPV in a level similar to iCaMKIIα (*p*=0.059; Figures 5Bii and 5C). By contrast, in PIGO(-)Szs iPV activity dropped below baseline immediately following seizure termination, remained relatively silent during the postictal period, and was significantly lower than postictal iPV activity of PIGO(+)Sz (*p*<0.05; Figure 5C).

While no elevated electrical activities were observed in bilateral hippocampi during the post-ictal period of PIGO(-)Szs, an increase in electrical activity was observed in the iCA1 and iDG regions in PIGO(+)Szs (*p*<0.01 and *p*<0.05, respectively in comparison to the same activities in PIGO(-)Sz). No increase in the EEG activity was found in the cCA1 or cDG in PIGO(+)Szs (Figures 5Bii and 5E), likely reflecting our previous findings showing a higher incidence of PIGOs in the iHIP (Figures 3Ai and 3Aii).

### PIGO(+)Szs showed delayed direct current potential shifts

Direct current (DC) potential shifts can be observed during various pathophysiological states such as cerebral hypoxia, spreading depolarization (SD), and epileptic seizures (40–43). To investigate the timing of the DC shifts relative to seizure termination, two additional PIGO(+) animals injected with pvIHKA were recorded with DC filtering disabled for spontaneous seizures (Figure 6). Among 99 seizures captured from these two animals, 82 were DC+, while 17 were DC-. Of the 82 DC+ seizures, 65 (79.3%) were PIGO(+)Szs, and 17 (20.7%) were PIGO(-)Szs. We observed that, while in PIGO(-)Szs, the DC deflections occurred, on average, prior to the seizure termination, in PIGO(+)Szs, the DC deflections occurred after seizure termination (*p*<0.0005; Figure 6B).

**Figure 6.**
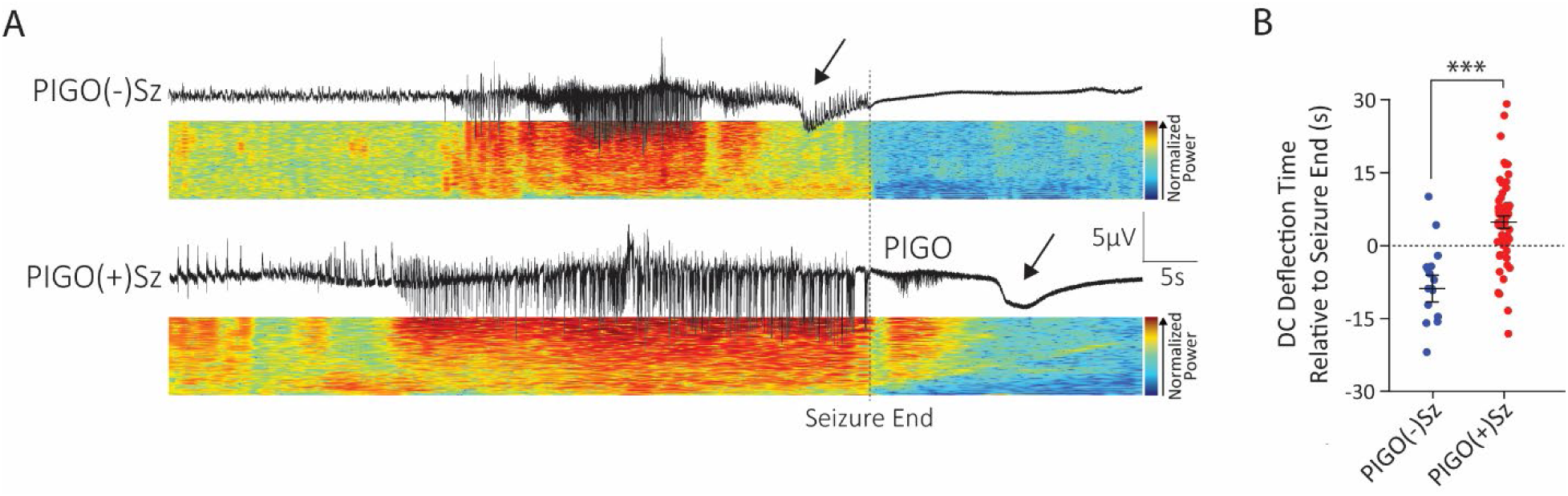
EEG direct current (DC) deflection with respect to PIGO. **A,** Sample EEG trace from spontaneous seizures recorded with DC-enabled electrodes. The upper trace illustrates a PIGO(-)Sz with a DC deflection occurring during the latter part of the ictal phase. The lower trace demonstrates a PIGO(+)Sz with a DC deflection in the post-ictal phase, coinciding with the presence of PIGO. Out of the 99 seizures captured, 82 seizures (82.8%) displayed DC shifts towards the end of the seizures, while 17 (17.2%) did not. Of the 82 seizures with DC shifts, 65 (79.3%) were PIGO(+)Szs, and 17 (20.7%) were PIGO(-)Szs. **B,** Time difference between seizure end (defined as y=0) and the occurrence of a DC deflection (n=82 seizures). In PIGO(+)Sz, the DC deflection was observed at an average of 4.39s (± 1.27 s, Median = 5.04 s) post-seizure end. In contrast, PIGO(-)Sz showed DC deflections occurring 8.75s ± 2.80s (Median = -7.92 s) before seizure end (*** P=0.00029; two-sample T-test).

### PIGO(+) animals had less degree of hippocampal sclerosis (HS)

The finding that pvIHKA animals, in which the recording sites were farther away from the IHKA injection, had more PIGOs led us to hypothesize that PIGOs may be associated with a reduced effect of the KA-induced excitotoxicity on the tissue at the recording sites. To test this hypothesis, histological analysis was performed to assess tissue changes in response to KA.

First, we visually inspected the hippocampal tissue surrounding the recording electrodes in the dorsal region of the iHip. Among 16 animals, 9 had a normal-appearing hippocampal structure at the recording region, while 7 animals showed HS with CA1 atrophy and DG enlargement (44) denoted as non-sclerotic recording region (NSRR) and sclerotic recording region (SRR) animals, respectively (Figure 7A). All 9 NSRR and only 1 SRR animals had PIGO(+)Szs (*p*<0.01; Figure 7B). NSRR animals exhibited a significantly higher proportion (*p*<0.01; Figure 7C) and incidence of PIGO(+)Szs compared to SRR animals (*p*<0.0005; Figure 7D).

**Figure 7.**
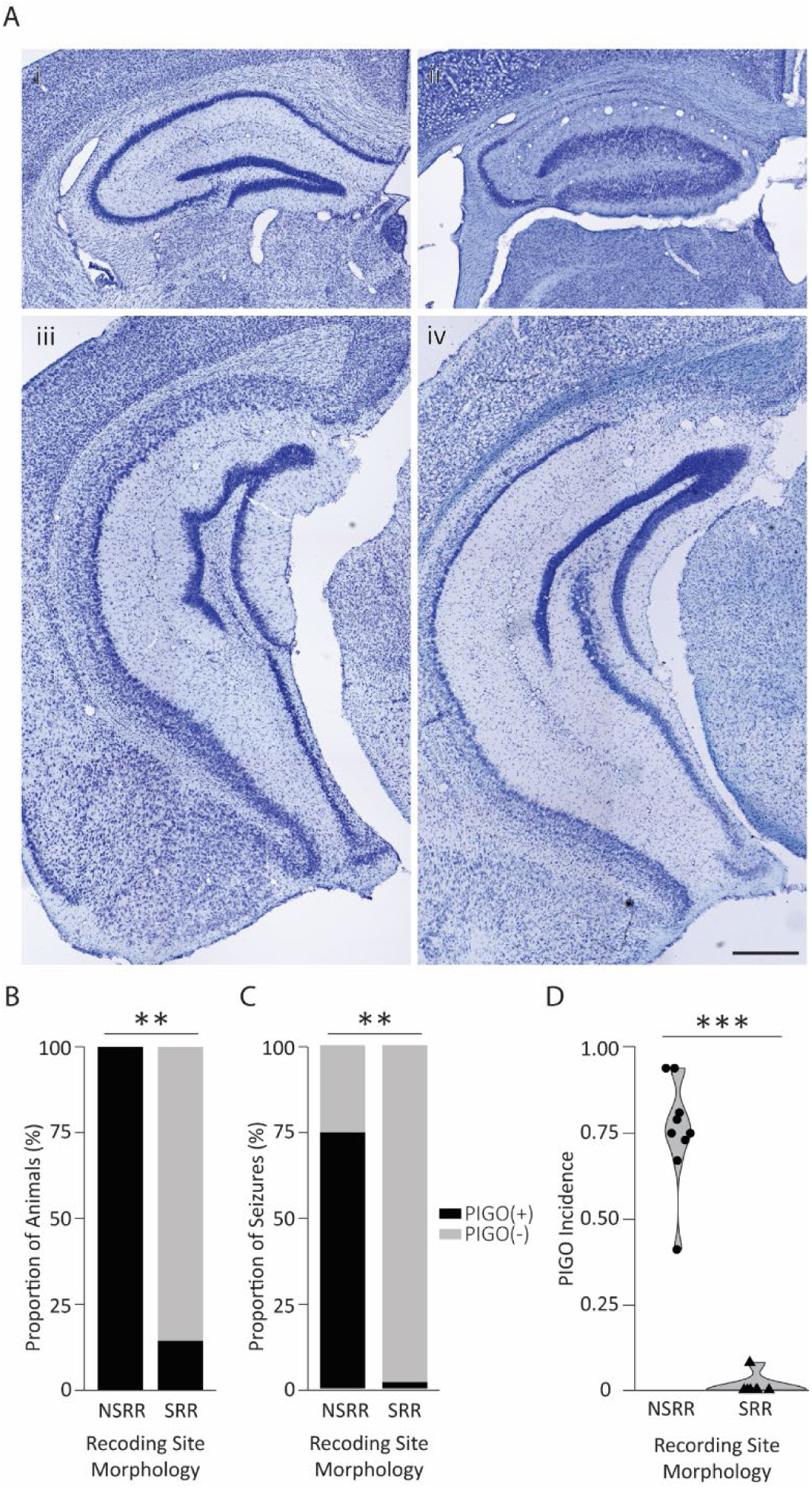
Hippocampal structural integrity and its influence on PIGO incidence. **A**, Representative Nissl-stained anterodorsal hippocampus section ipsilateral to the kainic acid injection of NSRR mice in **i** (n=9), and an anterodorsal hippocampus section ipsilateral to the kainic acid injection of SRR mice in **ii** (n=7). **iii** and **iv** represent posteroventral sections of the hippocampus from the same representative mice shown in **i** and **ii**, respectively. Scale Bar: 500µm. **B**, Proportion of animals in the NSRR and SRR groups that exhibited PIGOs following a seizure (9/9 animals from the NSRR group - 100%; 1/7 animals from the SRR group – 14%; **P<0.01; Fisher’s exact test). **C**, Cumulative proportion of PIGO(+)Sz in relation to the total number of seizures observed in each experimental group (NSRR group, n=9: 187/252 = 74.20%, SRR group, n=7: 2/164 = 1.21%, **P<0.01; Pearson’s Chi-squared test). **D**, PIGO incidence (number of PIGO(+)Sz / total number of seizures) for each animal from the NSRR and SRR groups. The NSRR group had a higher averaged PIGO incidence than the SRR group (75.44% ± 5.27 (range: 0.41 – 0.94) vs 1.14% ± 1.14 (range: 0 – 0.08), ***P<0.001; Wilcoxon rank sum test).

To expand on this analysis, we analyzed the structural characteristics of the hippocampus by measuring the thickness of bilateral CA1 and bilateral DG of PIGO(+) and PIGO(-) animals (Figure 8A; see Methods for details). Six additional animals that did not receive IHKA injections were also included for comparison (Naïve group). We observed that the hippocampi of PIGO(-) animals, on average, had more severe atrophy in the CA1 layer bilaterally than PIGO(+) (*p*<0.01), whose CA1 thickness was similar to the Naive animals (*p*=0.68; Figure 8B). This difference is likely due to a greater neuronal loss observed in the iCA1 layer of PIGO(-) animals in comparison to the same area of PIGO(+) animals (*p*<0.01). There was no difference in the CA1 thickness between the two hippocampi in either PIGO(+) or PIGO(-) animals (Supplementary Figure 3A).

**Figure 8.**
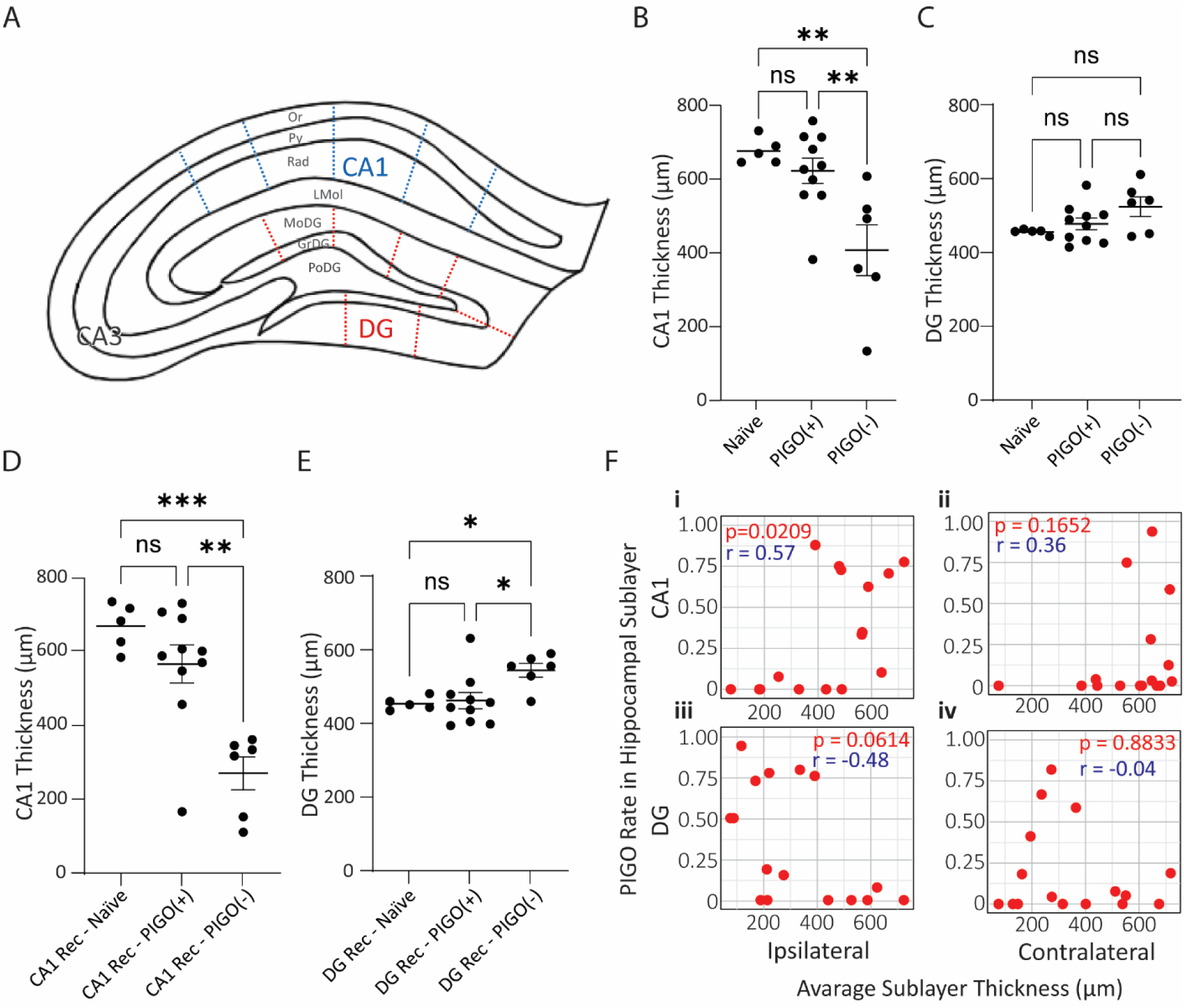
Quantification of the structural integrity of the hippocampus of mice injected with IHKA. **A**, Schematic of thickness quantification analysis of the hippocampal layers CA1 and DG in PIGO(+) and PIGO(-) animals. **B**, Average (± SEM) of the CA1 thickness of Naïve, PIGO(+), and PIGO(-) animals PIGO(-) animals had the thinnest CA1 layer in comparison to Naïve and PIGO(+) animals (CA1-Naive = 678.5 ± 15.9 µm, CA1-PIGO(+) = 624.6 ± 34.2 µm, CA1-PIGO(-) = 409.8 ± 68.8 µm). **C**, Average (± SEM) of the global DG thickness of Naive, PIGO(+) and PIGO(-) animals(DG-Naïve = 449.8 ± 3.42, DG-PIGO(+) = 471.6 ± 16.2 µm, DG-PIGO(-) = 518.6 ± 26.7 µm). **D**, Average (± SEM) of the CA1 thickness of Naive, PIGO(+) and PIGO(-) animals in recording areas. (CA1 Rec – Naive = 665.3 ± 27.9 µm, CA1 Rec – PIGO(+) = 563.2 ± 51.1 µm, CA1 Rec – PIGO(-) = 302.1 ± 57.6 µm). **E**, Average (± SEM) of the DG thickness of Naive, PIGO(+), and PIGO(-) animals in hippocampal Rec areas. (DG Rec – Naïve = 454.0 ± 7.8 µm, DG Rec – PIGO(+) = 462.1 ± 22.1 µm, DG Rec – PIGO(-) = 544.3 ± 18.8 µm P<0.05, Unpaired t-test). For **B**, **C**, **D** and **E**: ns P>0.05, * P<0.05, ** P<0.01, *** P<0.0005; One-Way ANOVA, Tukey’s multiple comparisons test as post hoc. Also, for **B**, **C**, **D** and **E**: Naïve animals (n=6), PIGO(+) animals: n=10, PIGO(-) animals: n=6. **F**, Correlation of sub-layer thickness and sub-layer PIGO incidence rates for each animal (n=16) analyzed with Spearman’s rank correlation coefficient.

We then performed a similar analysis on the structural integrity of the DG (Figure 8A). The averaged DG thickness across bilateral hippocampi was similar among PIGO(+) and PIGO(-) animals, and Naïve animals (*p*=0.08; Figure 8C). While the DG thickness was similar between the two hippocampi among PIGO(+) animals, the iDG was thicker than cDG in PIGO(-) animals (*p*<0.01; Supplementary Figure 3B).

We then repeated our analysis on the recording areas of the anterodorsal hippocampus, defined as brain slices with visible electrode tracks. In these areas, we observed significant differences in CA1 and DG thickness (Supplementary Figures 3C and 3D, respectively). While there was no difference in CA1 thickness between iCA1 and cCA1 within the recording areas of each group, the iCA1 of PIGO(-) animals was thinner than that of PIGO(+) and Naïve animals (p<0.0005 and p<0.0001, respectively). In contrast to the results observed in the CA1 layer, within PIGO(-) animals, iDG thickness was increased compared to cDG (p<0.01), as well as compared to iDG of PIGO(+) and Naïve animals (p<0.05). Non-recording areas of the hippocampus from PIGO(+) and PIGO(-) animals showed no thickness differences in either the CA1 or DG regions, neither between the two groups nor when compared to the same areas in Naïve animals (Supplementary Figures 3E and 3F, respectively). Taken together, our results suggest that, regardless of the IHKA injection site, PIGO(+) animals exhibit better-preserved hippocampal structural integrity compared to PIGO(-) animals. This difference is most pronounced at the sites of LFP recordings (Supplementary Figure 4).

To further support the finding above, we assessed the relationship between the PIGO incidence in each recording channel and the degree of HS in all animals (n=16). For this, the channel-specific PIGO incidence was correlated with the average thickness of the hippocampal layer for each animal (see Methods). A positive correlation was observed between the thickness of the iCA1 and the incidence of PIGOs. Specifically, animals exhibiting an iCA1 thickness larger than 500µm demonstrated elevated rates of PIGOs (*p*<0.05, Figure 8Fi). Although not significant, an inverse relationship between thickness and PIGO occurrence was observed in the iDG, where increased iDG thickness was associated with a reduced incidence of PIGOs (p=0.06; Figure 8Fiii).

### KA induces global astrogliosis in all IHKA animals regardless of the PIGOs emergence

Having shown that PIGO(+) animals have better-preserved hippocampal structural integrity compared to PIGO(-) animals, we next examine another important characteristic of HS: the proliferation and hypertrophy of astrocytes, also known as reactive astrogliosis (6–8). For this, we quantified the density and cell area of GFAP-positive cells in the hippocampus of PIGO(+) animals (n=7) and PIGO(-) animals (n=5) (See methods for details). We observed that animals from both PIGO(+) and PIGO(-) groups had similar increases in hippocampal astrocytic cell density (*p*<0.05; Figures 9A and 9B) and area (*p*<0.01; Figure 9C) relative to the hippocampi of Naïve animals (n=6). There was no difference in either astrocytic cell density or area between the two hippocampal hemispheres among all animals (Supplementary Figures 5A and 5B, respectively). In recording areas, the astrocytic cell density and area of both PIGO(+) and PIGO(-) animals were increased compared to those from the same regions of Naïve animals (Supplementary Figures 5C and 5D, respectively). However, the astrocytes in recording areas of PIGO(-) animals were larger than those in the same areas of the PIGO(+) animals (*p*<0.05; Supplementary Figure 5D).

**Figure 9.**
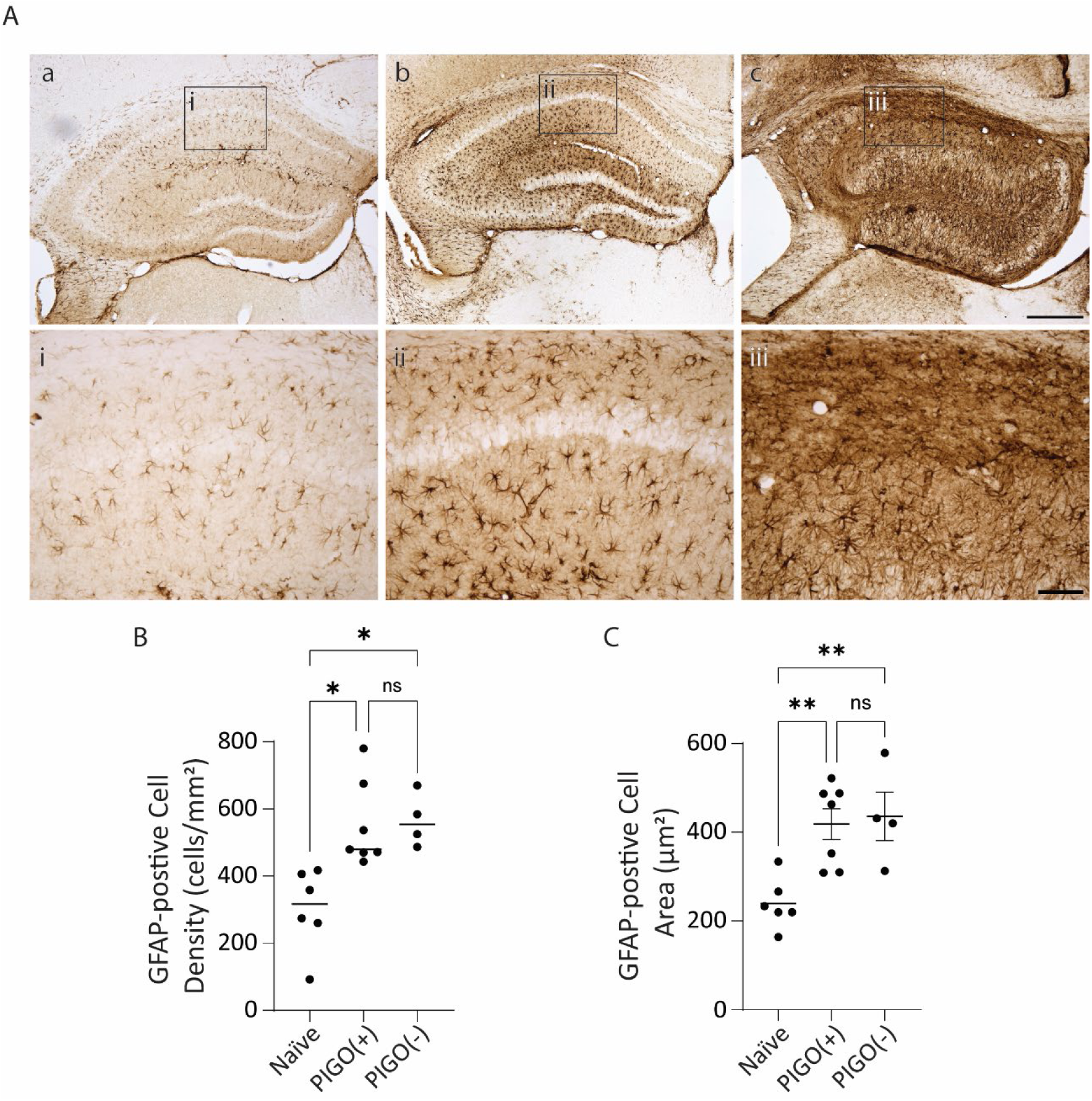
Kainic acid-induced global astrogliosis in both PIGO(+) and PIGO (-) animals. **A,** Representative GFAP-stained dorsal hippocampus sections of Naïve, PIGO(+) and PIGO(-) mice. Subfigures **a, b,** and **c** display antero-dorsal hippocampal sections of a Naïve, PIGO(+), and PIGO(-) mice, respectively. Insets show magnified micrographs of GFAP-positive cells in the hippocampal CA1 sublayer indicated by boxes **i**, **ii**, and **iii**. Scale Bars: **a**-**c**: 500µm, insets i – iii: 100µm. **B**, Average density of GFAP-positive cells for each experimental group (Mean ± SEM). (Naïve = 305.6 ± 47.4 cells/mm^2^, PIGO(+) = 507.6 ± 58.0 cells/mm2, PIGO(-) = 546.1 ± 44.8 cells/mm2; Naïve vs. PIGO(+): P<0.05, Naïve vs. PIGO(-): P<0.05; PIGO(+) vs. PIGO(-): P=0.94; One-way ANOVA; Tukey’s multiple comparisons as a *post hoc* test). **C**, Average area of GFAP-positive cells (in µm^2^) measured from the single astrocytes counted in each hippocampal section of Naïve, PIGO(+), and PIGO(-) animals (Naïve = 239.4 ± 23.2 µm^2^, PIGO(+) = 418.6 ± 34.6 µm^2^, PIGO(-) = 435.8 ± 54.6 µm^2^; Naïve vs. PIGO(+): P<0.01, Naïve vs. PIGO(-): P<0.01, PIGO(+) vs. PIGO(-): P=0.89; One-way ANOVA; Tukey’s multiple comparison as a *post hoc* test).

### PIGOs were observed in human patients with unilateral mTLE

To demonstrate the relevance of the experimental findings on PIGOs to human patients with mTLE, we reviewed data from two adult patients who underwent intracranial EEG evaluation. Two patients with unilateral mTLE were selected for this analysis. The first patient was a 46-year-old right-handed male who experienced epilepsy onset at age eight and had between two and four focal impaired awareness seizures per month, despite undergoing multiple anti-seizure medication trials. Non-invasive EEG captured five right temporal onset seizures, but structural 3T MRI revealed no epileptogenic lesions, including no mesial temporal sclerosis, focal cortical dysplasia, or space-occupying lesions (Figure 10Ai). FDG-PET showed no distinct areas of focal hypometabolism to suggest an epileptogenic focus (Figure 10Aii). Due to his non-lesional MRI and PET findings, he underwent stereo-EEG (sEEG) monitoring with 15 electrodes on the right hemisphere to delineate the SOZ. Five seizures were captured during the intracranial monitoring period. Our analysis revealed four of these five seizures had at least one hippocampal recording channel exhibiting a PIGO. In one seizure shown in Figure 10Aiii, the recording channel located in the anterior hippocampus exhibited increased EEG power in the gamma band that began approximately 45 seconds after the seizure termination and lasted for approximately 20 seconds. These EEG recordings confirmed the SOZ in the right mesial temporal structures, and he subsequently underwent laser interstitial thermocoagulation of the right amygdala and hippocampus, remaining seizure-free for eight years to date.

**Figure 10.**
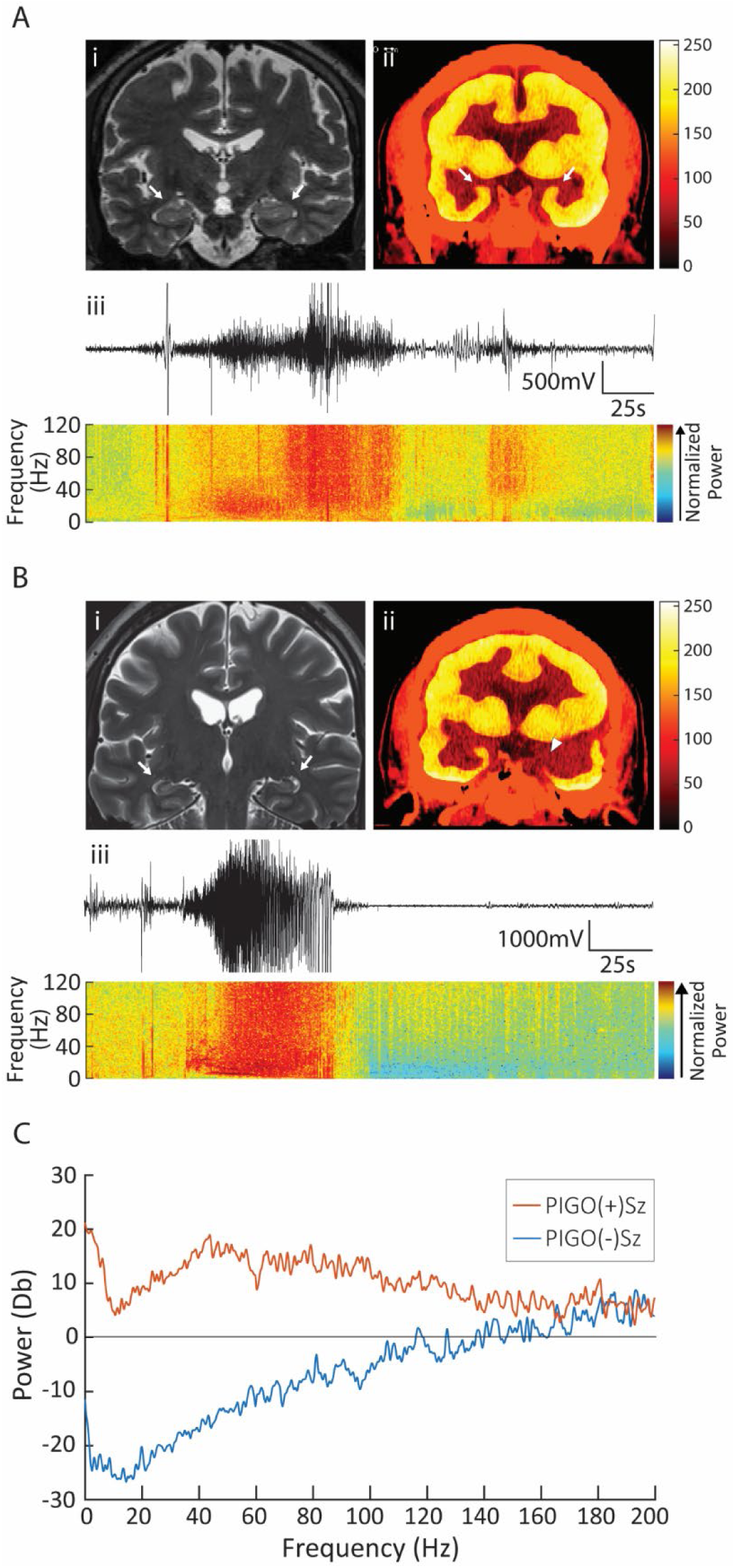
Evidence of PIGO in a patient with mTLE. **A,** PIGO(+) patient coronal T3-MRI slice (**i**) and FDG-PET image (**ii**). Note that both the MRI and FDG-PET scan resulted in structural and metabolic symmetry between both hippocampal lobes (arrows). **iii** shows a time-synchronized sEEG and spectrogram of a seizure event recorded in the right hippocampus. **B,** PIGO(-) patient T3-MRI slice (**i**), showing structural symmetry between the hippocampal lobes (arrows). The FDG-PET scan of this patient (**ii**) revealed hypometabolism in left temporal structures (arrowhead). **iii** shows a time-synchronized sEEG and spectrogram of the PIGO(-)Sz obtained from an intracranial left-hippocampal recording of a seizure event. **C,** Comparison of the power spectrum in the post-ictal period between the PIGO(+)Sz showed in **Aiii**, and the PIGO(-)Sz showed in **Biii**.

As a comparison, the second patient was a 53-year-old right-handed male with epilepsy onset at age 39. He experienced one focal impaired awareness seizure per week, despite multiple trials of anti-seizure medications. Non-invasive EEG captured four habitual seizures with left temporal onset, and structural 3T MRI was unremarkable (Figure 10Bi). However, FDG-PET revealed marked hypometabolism in the left mesial temporal region, which was prominently asymmetric compared to the contralateral side (Figure 10Bii). Given normal neuropsychological performance, ictal involvement of the dominant temporal region, and a non-lesioned MRI, the patient underwent sEEG to precisely localize the SOZ for a potentially more selective destructive procedure aimed at preserving cognitive function. Two seizures were captured during his invasive monitoring; neither exhibited PIGOs in the post-ictal period at any hippocampal electrode (Figure 10Biii). Intracranial sEEG confirmed the SOZ in the left medial temporal structures, and the patient subsequently underwent laser interstitial thermocoagulation of the left amygdala and hippocampus. This patient did not achieve seizure freedom despite the destruction of left mesial temporal lobe structures.

Next, we calculated the power spectrum from post-ictal activity of the two patients’ seizures (Figure 10C). In the PIGO(+)Sz, we observed an increase in power starting around 20 Hz, with a peak at 40 Hz. The increased power extended from 40 to 100 Hz and gradually tapered off in the hyper-gamma range above 100 Hz. In contrast, the PIGO(-)Sz showed decreased power in frequencies below 140 Hz, with a slight elevation in the hyper-gamma range, overlapping with the PIGO(+) power spectrum. This frequency signature was consistent with the gamma frequency elevation observed in the animal data.

## Discussion

In this paper, we report a distinct EEG activity, post-ictal gamma oscillation (PIGO), observed following spontaneous seizures in a specific phenotype of IHKA-induced mice with mTLE. This phenotype exhibits a less severe degree of hippocampal sclerosis (HS) at the recording site and in the hippocampus overall, characterized by reduced CA1 atrophy and decreased DG granular cell dispersion (Figure 8), as well as less astrogliosis (Figure 9). PIGOs are largely absent in a phenotype with extensive HS (Figure 2). PIGO is characterized by peak spectral activity in the gamma frequency, which is traditionally thought to be driven by activity from PV-INs in the hippocampus (45). Fiber photometry recordings revealed a large post-ictal surge in iPV photometry during PIGOs in the same animals, consistent with the involvement of PV-INs in this electrical activity (Figure 5).

PIGO has been reported by other studies conducted in animals (46) and in epileptic patients (47). Specifically, a low amplitude, 40-80 Hz gamma-range activity was observed by Bragin et al. (1997) as a response to the stimulation of the perforant path or the commissural system of the hippocampus of rats, termed “afterdischarge termination oscillation” (ATO). Similar to our results, ATOs were observed initially in CA1, eventually appearing in other hippocampal regions in a delayed fashion (46). Their findings suggested that PV-INs do not play a significant role in the generation of ATOs, since PV-INs showed little or no activity during ATOs. These fast-spiking GABAergic neurons are known to provide feedforward and feedback synaptic inhibition and prevent excessive excitation in neural networks (48). However, dysfunctional PV-INs are described to play a role in the generation and maintenance of hypersynchronous ictal activity (49–51). In the current study, in addition to being associated with the hypersynchronous activity during the ictal period in PIGO(+)Szs and PIGO(-)Szs, rises in PV-IN [Ca^2+^]_i_ activity were associated with PIGO activity in iCA1 (Figure 5B-D). A possible underlying mechanism for PIGOs may lie in an interneuron network model. Interestingly, even if PIGOs were present at bilateral CA1 and DG on the EEG, changes in Ca^2+^ activity were only detectable in the iHip (Figure 5B). Previous studies have shown that gamma oscillations depend on GABA_A_-receptor-mediated inhibition, suggesting that these oscillations are primarily generated by networks of inhibitory interneurons (45, 52). Fast-spiking parvalbumin-expressing basket cells are key components of the hippocampal interneuron network. Based on mutual inhibition, this network, assuming slow, weak, and hyperpolarizing synapses, can generate synchronized gamma activity when exposed to a tonic excitatory drive such as generalized seizures.

The increase in [Ca^2+^]_i_ among both pyramidal cells and PV-INs suggest that PIGOs may be associated with spreading depolarization (SD), a phenomenon known to follow the end of seizures (53). The increase of [Ca^2+^]_i_ accompanied by diminishing EEG activity in our recording (Figure 5) is characteristic of an SD-induced depolarization event. The timing of these events also corresponds to a large electrophysiological direct current (DC) deflection (Figure 6), characteristic of increased extracellular K^+^ during SD (54). Since SD depolarizes all neuron types indiscriminately, the larger iPV photometry response indicates a larger surviving PV-IN population in the mice phenotype exhibiting PIGO. We, therefore, suspect that PIGO captures the activity driven by the surviving hippocampal PV-INs, emerging near or after the end of the seizure and prior to a delayed SD onset in IHKA epileptic mice with mild HS.

In patients, post-ictal gamma-range activity (>25Hz) was observed by Bateman et al. in intracranial sEEG recordings of 16 epileptic patients with focal to bilateral tonic-clonic seizures (FBTCs) (55). These patients had the seizure onset zone predominantly in the temporal lobe, with a few in the frontal, parietal, and occipital lobes. The elevated gamma oscillations observed following the seizure termination were associated with marked low-frequency attenuation, thus named “intracranial post-ictal attenuation” (IPA) (55). This finding is similar to our observation that PIGOs display elevated gamma-range activity and attenuation of low-frequency range activity (Figure 1B and Figure 3F). Additionally, the spectral characteristics of IPA in our human recordings were associated with attenuated signals relative to the ictal phase and increased gamma activity relative to the preictal period. Interestingly, IPA was timely associated with a wide range of behavioral manifestations, including flaccid paralysis. In contrast, PIGOs in our study were associated with a near absence of any motor symptoms. The authors hypothesized that IPA resulted from ongoing seizure-related neuronal activity in unrecorded brain regions. Additionally, they speculated that subcortical, possibly brainstem structures, drive IPA. However, we believe that PIGOs are unlikely to be a continuation of ongoing ictal activity, given their distinct spectral characteristics compared to the preceding ictal period (Figure 3F).

One of the key findings of the present study is that PIGOs were only observed in a subgroup of animals. Our results indicated that the emergence of PIGOs predicted less structural alteration in response to the KA (Figures 7, 8, and Supplementary Figure 3). To the best of our knowledge, this is the first report of an electrophysiological biomarker for underlying tissue pathology in focal epilepsy. Due to the limited spatial sampling in the current study, it is possible that PIGO(-) animals could exhibit PIGOs in regions of the bilateral hippocampi that were not covered by electrodes, particularly in areas with a lower degree of HS. However, in the PIGO(-) animals, PIGOs were not recorded in either hippocampus, even though the hippocampus contralateral to the site of IHKA injection was observed to have a lesser degree of HS (Figure 8 and Supplementary Figure 4). Furthermore, our histological analysis suggested that the PIGO(+) animals had an overall lower degree of HS across bilateral hippocampi compared to the PIGO(-) animals (Figure 8F). Indeed, the ATOs reported by Bragin et al. appeared in response to electrical stimulation in healthy wild-type rats with normal hippocampi (46), further supporting the notion that PIGOs were associated with animals that have a lesser degree of HS.

The postictal periods containing PIGOs exhibit more elevated electrical and cellular activities than the postictal periods of seizures absent of PIGOs (Figure 5). Seizure termination is commonly followed by a phenomenon characterized by prolonged post-ictal generalized electroencephalographic suppression (PGES) (27, 28, 30). In our study, not all animals and patients with mTLE have seizures followed by PIGOs. If PIGOs in the postictal period displace PGES, PIGO(+) animals and patients may be less prone to developing PGES. There is an association between PGES and impaired consciousness, cardiorespiratory arrest, and even sudden unexpected death in epilepsy (SUDEP) (56–58). Although the mechanism of PGES is poorly understood, the leading hypothesis is that subcortical thalamic nuclei, which are implicated in critical arousal processes, are deactivated following a generalized seizure, altering the network mechanism that drives the widespread postictal EEG changes (27, 57, 59, 60). Moreover, deep subcortical nuclei, such as thalamic and brainstem structures connected to cortical brain regions, are disrupted after a generalized seizure (57, 59–61). Since PIGO(-) animals and patients may be more prone to PGES, their epileptic network may have a higher likelihood of involving deep nuclei. Existing evidence suggests that the involvement of subcortical structures in the epileptogenic network is a risk factor for postoperative seizure recurrence (62, 63).

Finally, as a proof-of-concept for the translational potential of PIGOs, we selected two patients with mTLE who underwent intracranial sEEG recordings for SOZ localization. Both patients had mTLE, but one had PIGO(+)Szs, while the other did not (Figure 10). While both patients had non-lesional, normal-appearing MRIs, the PIGO(-) patient showed markedly reduced metabolism in the left mesial temporal region on FDG-PET. The mesial temporal region was confirmed by sEEG as the SOZ for both patients. Although no direct histological comparison can be made between the PIGO(+) and PIGO(-) patients because the surgical intervention (laser ablation) does not retrieve brain tissue, the difference in the FDG-PET studies between the two patients may suggest that PIGO(-) patients have a more severe form of HS (64, 65). This observation is an important step in understanding whether this electrophysiological biomarker applies to mTLE patients. Moreover, after both patients underwent surgical treatment, the PIGO(+) patient responded better to the intervention than the PIGO(-) patient. This difference in treatment outcomes is consistent with our hypothesis that PIGO is a biomarker for mTLE with a lesser degree of HS and an epileptic brain network with less involvement of subcortical structures.

## Methods

### Experimental Animals

In this study, fifteen male and nine female mice were used. Thus, sex was not considered as a variable. The experiments were conducted in adult (8-10 weeks old) male and female inbred homozygous PV-Cre knock-in mice (PV-cre^+/+^, B6.129P2-Pvalb<tm1(cre)Arbr>/J, Strain #: 017320 – The Jackson Laboratory). Mice were housed with their littermates (1-4 mice per cage) until the surgical procedures, with environmental enrichment conditions and food and water provided *ad libitum* under standardized temperature, humidity, and a 12 h light/dark cycle (lights on from 6:00 am to 6:00 pm).

### Animal Surgeries

Four to six-month-old PV-cre^+/+^ mice underwent intra-hippocampal kainic acid injection and electrode implantation surgery. Mice were placed in a chamber and anesthesia was induced with isoflurane (∼2.5%). The animals were transferred to a heating pad and kept warm while isoflurane-induced anesthesia was maintained throughout the whole surgical procedure (∼1.5% during maintenance). Sustained release buprenorphine (EthiqaXR, 3.25 mg/kg, s.c.) and a local anesthetic (Bupivacaine, 2.5mg/kg, s.c.) was administered to alleviate pain and discomfort. The skull was exposed and through burr holes made by a micromotor drill (Stoelting Co., Wood Dale, IL, USA). 50, 75, or 100 nL of kainic acid (KA, 20mM; HelloBio, Princeton, NJ, USA; Supplementary Figure 6) was stereotaxically microinjected unilaterally in the CA1 layer of the dorsal hippocampus (adIHKA group, coordinates from bregma: anteroposterior (AP) = -2.06 mm, mediolateral (ML) = -1.6 mm, and dorsoventral (DV) = -1.37 mm, Supplementary Figure 6); or with 50nL (20mM) of KA in the CA1 layer of the ventral hippocampus (pvIHKA, AP = -3.64, ML = -3.2 mm, DV = -3.3 mm), at a rate of 25nL/min using a Hamilton syringe (Neuros 7001, Hamilton Co., Reno, NV, USA) attached to an automatized injector (Quintessential Stereotaxic Injector, Stoelting Co., Wood Dale, IL, USA). After the injection, the needle was maintained in place for five minutes to avoid backflow and then slowly retracted. In the same surgery, the animals were implanted with a 5-channel electrode ensemble (See Figure 1A), composed of four recording electrodes (F12146, P1 Technologies, Roanoke, VA, USA) positioned at bilateral CA1 (AP, -2.06 mm; ML, ±1.6 mm; DV, -1.4 mm) and DG (AP, -2.06 mm; ML, ±1.6 mm; DV, -1.9 mm), with the remaining electrode connected as reference. The electrodes were ensembled in a connector pole (Mini6, Plastics One, Roanoke, VA, USA) that served as an interface between the electrodes and the recording system and was secured to the skull using dental acrylic cement. After surgery, the animals were individually housed in a clean cage on a heating pad to recover. Two hours after the KA injection, animals were administered two to three systemic low-dose Diazepam injections (5mg/kg, i.p.) every 30 minutes to halt *status epilepticus* (SE) and reduce mortality.

### EEG Recordings

After a recovery time of at least three days after the IHKA injection, each animal was placed in the recording cage with food and water *ad libitum* and environmental enrichment, and a cable was connected to the electroencephalography (EEG) headset ensemble for video-EEG recordings. In addition to a cable for LFP recordings, the pvIHKA animals also had an optical patch cord connected to each implanted optic fiber (400μm, 0.57NA, MFC_400/430-0.66_4mm_ZF1.25(G)_FLT, Doric Lenses). EEG signals were collected at 3 kHz, amplified using a TDT RZ10x or RZ5 system (Tucker-Davis Technologies, Alachua, FL, USA), band-pass filtered from 3 to 1000 Hz, and notch filtered to attenuate 60Hz noise and its harmonics. EEG traces were manually reviewed offline using a custom-made MATLAB application (MathWorks), where the relative times of seizure onset and termination, and PIGO onset, termination, and specific channel location were marked. Seizure onset was defined as the earliest change from baseline detectable prior to the generalized ictal activity. The severity of the seizures was assessed offline by the analysis of the videos recorded simultaneously with the EEG. The seizure severity of the seizures was scored using a modified Racine scale (RS) adapted from Erun *et. al.* (2019) (37) as follows: (0), no behavioral alteration; (1), arrest or sudden motion; (2), mouth chewing and/or head nodding; (3), forelimb clonus; (4), fore- and hindlimb clonus and rearing; (5) loss of posture/falling; (6), brief wild running or jumps; (7), severe wild running or jumps. For seizures with PIGOs present (PIGO(+)Sz), a RS was assigned for the period during which PIGO was present on the EEG.

### PIGO Identification and Incidence Calculation

Each animal’s seizures were classified as either PIGO-positive (PIGO (+)) or PIGO-negative (PIGO (-)) by two independent raters based on the presence of a PIGO following a seizure. PIGO incidence was calculated as the percentage of total seizures that were PIGO(+). Cumulative calculations were conducted by grouping all seizures within an animal category. Channel-specific PIGO incidence was calculated by tagging each channel of a seizure as PIGO(+) or PIGO(-). The channel-specific PIGO incidence was then calculated by dividing the number of PIGO(+) channels (i.e., the number of times a channel had a PIGO following a seizure) by the total seizure count.

### Principal Component Analysis

Principal component analysis (PCA) was performed on the PIGO channel combinations to identify patterns in PIGO emergence across recording channels. The data matrix was standardized, and PCA was applied to reduce the dimensionality and extract the principal components. K-means clustering was then conducted on the principal component scores to classify the patterns into three distinct clusters.

### Frequency Analysis

Power spectrums were calculated using MATLAB’s Signal Processing Toolbox with a Discrete Fourier Transform method with frequency limits from 0Hz to 150Hz and a frequency resolution of 3Hz for both the PIGO phase and ictal phase for each channel with a PIGO. A baseline window 20 seconds prior to seizure onset was subtracted from PIGO and ictal power spectrums to minimize noise and normalize the power spectrums. The peak frequency, defined as the frequency of maximum power contributing to the EEG signal, was calculated for each EEG channel and tabulated.

### PIGO Correlation Analysis

Pearson’s correlation coefficients were calculated to assess EEG signal during the PIGO phase of PIGO(+)Sz and PIGO(-)Sz. In PIGO(+)Sz, the PIGO phase was defined by time segment where a PIGO was present. When PIGOs were present in multiple channels, the overlapping PIGO time segment was used for all correlations between channels. For PIGO(-)Sz , the PIGO phase was defined by the average PIGO duration and average PIGO onset relative to the seizure end. Therefore, for each individual seizure, correlations were calculated using time segments of equal duration across all four channels. The correlation coefficients were then Fisher Z-transformed, and a 20-second baseline correlation was subtracted to exclude electrical variances in EEG signals.

### Histology Processing

Upon the completion of recording experiments, mice were deeply anesthetized with isoflurane and then transcardially perfused with ice-cold 4% paraformaldehyde (PFA) in 1X phosphate-saline buffer (PBS – pH7.4). Extracted brains were post-fixed in 4% PFA and then transferred to 30% sucrose solution in 4% PFA until being sectioned in coronal sections (30μm) in a cryostat (Leica CM3050S, Leica Microsystems, Wetzlar, Germany) and were stored in cryoprotectant at -20°C until being processed. Brain slices obtained from Naïve animals were used as a control for histological analysis.

To assess CA1 atrophy and granular cell dispersion (GCD) in the animals treated with IHKA and Naïve animals, six coronal brain sections of each animal were selected and processed using Nissl staining. The selection of the sections was made with a uniform and similar caudo-rostral distribution among the animals to cover the whole anteroposterior extent of the hippocampus. AP distance of the slices from Bregma were approximated at the following slice intervals: Slice 1: 1.30–1.65 mm, Slice 2: 1.65–2.00 mm, Slice 3: 2.00–2.35 mm, Slice 4: 2.35–2.70 mm, Slice 5: 2.70–3.10 mm, and Slice 6: 3.10–3.50 mm. Briefly, the brain sections were mounted on gelatin-coated slides, dried, rehydrated in a series of alcohol baths, stained in 0.5% cresyl violet, washed, dehydrated in increasing ethanol gradient, cleared in xylene, and coverslipped using Permount mounting medium (Fischer Scientific, Hampton, NH, USA). Hippocampal images were taken using a brightfield Leica microscope (Leica DM 4B; Leica Microsystems, Wetzlar – Germany). CA1 atrophy and/or GCD in the hippocampus of the epileptic mice were characterized by two different approaches: (i) visual inspection of the brain slices by a trained person, (ii) measurement of the thickness of the hippocampal CA1 layer and the DG using ImageJ (see Figure 6A for schematic of the measurements; ImageJ, U.S. National Institutes of Health, Bethesda, MD, USA). For CA1 thickness assessment, 5 equidistant measurements of the CA1 layer (stratum oriens, stratum pyramidale, and stratum radiale) of each hippocampal lobe were made in regular length intervals, and averaged. For DG structural integrity analysis, we analyzed the thickness of the granular cell layer (GCL – stratum granulosum) and the molecular layer (ML, outer, middle, and inner ML) of the supra- and infrapyramidal blades of the DG. To analyze DG thickness, 7 equidistant measurements were made, and averaged. The thickness measurements from CA1 and DG were collected bilaterally in each brain slice.

To assess astrogliosis in the hippocampus of adIHKA and pvIHKA animals, six brain slices from each animal (as in for Nissl staining) were processed for glial fibrillary acidic protein (GFAP) immunohistochemistry (IHC). Free-floating coronal brain sections were washed 5×5min in tris-buffered saline (TBS) containing 0.05% Triton X-100 (TBST) and incubated in 0.3% hydrogen peroxide for 30 minutes. The slices were washed in TBST 3×5 min and blocked for 60 minutes at room temperature in goat blocking buffer (GBB). Sections were then incubated overnight at 4°C in GBB containing the primary antibody rabbit anti-GFAP (MA5-35237, Invitrogen, Waltham, MA, USA). The sections were then washed 3×10min in TBST and incubated for 90 minutes at room temperature in GBB containing the biotin-conjugated goat anti-rabbit secondary antibody (SAB4600006, Sigma-Aldrich, San Luis, MI, USA). After 3 × 10 min washes in TBST, the slices were incubated in avidin-biotin complex (ABC) solution (Vector Laboratories, Newark, CA, USA), washed 3×5 min with TBST, and incubated in 3,3’-Diaminobenzidine (DAB) substrate solution (Vector Laboratories, Newark, CA, USA) for up to 10 min until the reaction product was visualized. After washing 3×5 min in TBST, the slices were mounted on slides and then allowed to dry overnight. Sections were dehydrated in a series of alcohol baths, cleared in xylene, and cover-slipped with Permount-G mounting medium (SP15-100 UN1294; Fisher Scientific, Waltham, MA, USA). Bright-field images were acquired using an image scanning function at 10x magnification on a brightfield Leica microscope (Leica DM 4B, Leica Microsystems, Wetzlar, Germany). Cell counting and astrocytic cell area measurements were performed using the particle analyzer function of ImageJ (ImageJ, U.S. National Institutes of Health, Bethesda, MD, USA).

### Ca^2+^ Fiber Photometry analysis of Principal neuron and PV-IN activity in the hippocampus

For the analysis of Ca^2+^ fiber photometry activity in glutamatergic neurons and parvalbumin-positive interneurons (PV-INs), the mice underwent a first stereotaxic surgery where they were microinjected with an adeno-associated viral vector (AAV) cocktail containing AAV8-CaMKIIα-jRCaMP1b, targeting principal neurons (CaMKIIα-RCaMP, 100nL – Stanford Vector Core) and the cre-dependent pGP-AAV-syn-FLEX-jGCaMP8m-WPRE (AAV9), targeting PV-INs expressing cre recombinase (PV-GCaMP, 150nL – Addgene #162378-AAV9 (66)) in the CA1 area of the dorsal hippocampus (coordinates: AP, -2.06 mm; ML, ±1.6 mm; DV, -1.4 mm) at least 21 days before the second stereotaxic surgery for pvIHKA injection and electrode implantation. In this second surgery, in addition to the six electrodes for local field potential (LFP) recording as in the animals from the adIHKA group, mice from the pvIHKA group were implanted with two optic fibers (0.66 NA, 400 mm diameter, Doric Lenses, Quebec – Canada). The optic fibers were positioned above the CA1 layer bilaterally in the dorsal hippocampus (AP, -2.06 mm; ML, ±1.6 mm; DV, -1.1 mm) for Ca^2+^ fiber photometry assessment.

The cellular activity of principal neurons and PV-INs, recorded simultaneously with video-EEG, was captured through the two optic fiber cannulas implanted. Ca^2+^ fiber photometry signals were recorded by capturing the fluorescence deflections (ΔF/F) in response to calcium binding into the genetically encoded calcium indicators (GECIs) RCaMP and GCaMP. Fiber photometry acquisition was performed interchangeably between two TDT-based systems. The RZ10x system has inbuilt LEDs and photosensors, and they were coupled to two passive Doric FMC6 mini cubes with optical filters only (FMC6_IE(400-410)_E1(460-490)_F1(500-540)_E2(555-570)_F2(580-680)_S, Doric). The second system is RZ5-based coupled to two ilFMC6 Doric mini cubes with inbuilt LED and photosensors (ilFMC6-G3_IE(400-410)_E1(460-490)_F1(500-540)_E2(555-570)_F2(580-680)_S, Doric) and a single 4-port LED driver (LEDD_4, Doric) that controls both ilFMC6 mini cubes. Photometry signals from RCaMP and GCaMP were collected simultaneously, excited at 560nm and 465nm respectively, with an isosbestic signal excited at 405nm, illuminated using corresponding TDT’s Lux LEDs when recording with the Rz10x system or ilFMC6 inbuild LEDs when recording with the RZ5 system. Doric mini cubes combined all three excitation lights and collected the emission fluorescence from a single patch-optical chord for each hippocampus cannula. Each animal, therefore, was connected to two patch cords and two mini cubes, one for each side of the hippocampus. The lock-in amplification approach was adopted for fiber photometry signal collection, using modulation frequencies 211Hz, 330Hz, and 530Hz to separate the three photometry channels. Modulation and demodulation were controlled by the TDT’s control software. Photometry signals were collected at 6kHz, demodulated down to 1kHz, and low-pass filtered at 3Hz for storage.

Four Ca^2+^ fiber photometry signals were visualized and plotted using a custom-made MATLAB application (MathWorks). For visualization and data extraction, we scaled the ictal and post-ictal periods, so seizures and PIGOs with different durations could be stacked into equal-duration data blocks: (i) pre-ictal period (baseline, -60 to -0s), (ii) ictal period (0s to 60) and (iii) post-ictal period (60 to 120s). ΔF/F and coastline index data were extracted in 1-second bins from each seizure using the same custom-made MATLAB application (MathWorks) as for the visualization. To calculate the levels of [Ca^2+^]_i_ or CI for each animal, the ΔF/F 1-second bins and coastline index values were averaged across each data block from each seizure and then averaged across the seizures. [Ca^2+^]_i_ signal peak timings were identified by the identification 1-second bin correspondent to the maximum ΔF/F value for the ictal or post-ictal data blocks of each seizure.

### EEG Coastline Index Calculation

The levels of EEG activity in each hippocampal recording channel were quantified by calculating the coastline index (CI) for posterior correlation analysis with Fiber Photometry signals. This computational method was shown to be sensitive to quick and high fluctuations of ictal activities (38, 39) by measuring the size of the EEG voltage change from sample to sample per unit of time (expressed in V/s; see algorithm below).

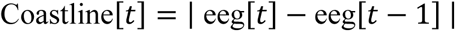

The CI for each electrode was baseline-adjusted such that the average baseline coastline from t=-60s to 0s was set to 0 V/s; subsequently, the negative adjusted coastline index was reset to 0 V/s. CI signals were visualized, plotted, and extracted using a custom-made MATLAB application (MathWorks) using the same time-windows as for Fiber Photometry signals.

### Human data

Two illustrative patients with suspected unilateral temporal lobe epilepsy diagnosed based on International League Against Epilepsy criteria (67) underwent stereo-EEG to help confirm the seizure onset zone for possible surgical intervention. The location of intracranial recording electrodes was determined by co-registering the electrode positions in post-surgical CT with pre-surgical MRI T1-weighted imaging using open-source electrode reconstruction pipelines (68).

Spontaneous ictal events were analyzed. A 60Hz notch filter was implemented to remove electrical noise, and a 0.5Hz-500Hz Butterworth bandpass filter was implemented to remove sub-0.5Hz noise. Spectrograms of the seizure EEGs were calculated with Short-Time Fourier Transformation with a Hamming window length of 1000. Power spectra of post-ictal windows were normalized by subtracting a baseline window of the same duration prior to seizure onset. For PIGO(-) seizures, a post-ictal comparison window of the same duration and onset relative to seizure resolution was used for power spectrum calculations.

### Statistical Analysis

All statistical analyses were conducted using RStudio Software (Version 2023.03.0; Boston, MA, USA) or GraphPad Prism 8 (Dotmatics, Boston, MA, USA). Parametric or nonparametric tests were selected based on data distribution and used to assess significance (p < 0.05). For all results, the significance threshold was placed at α = 0.05, and corrections for multiple comparisons are reflected in P values. To compare 2 independent groups of values with respect to a numeric outcome, t-tests were used. For multiple comparisons between variables, Two-Way Analysis of Variance (ANOVA) was employed, followed by post hoc pairwise comparisons using the Tukey test. To assess linear relationships between variables. Pearson or Spearman correlation analyses were applied. To evaluate differences in binomial counts between groups when the sample size was adequate, Pearson’s Chi-squared test and Fisher’s exact test was utilized for small sample sizes. The Wilcoxon rank-sum test was also employed to compare the distribution of values between groups. Results are presented as mean ± standard error of the mean (SEM) unless otherwise specified.

## Animal study approval

All animal procedures were conducted in accordance with approved Rutgers Institutional Animal Care and Use Committee (IACUC) protocols within an Association for Assessment and Accreditation of Laboratory Animal Care (AAALAC) accredited facility.

## Human study approval

Human data were collected following Emory University Institutional Review Board (IRB) approval for retrospective collection of clinically acquired data.

## Author Contributions

DV, designing research studies, conducting experiments, acquiring data, analyzing data, writing of the manuscript; FT, designing research studies, conducting experiments, acquiring data, analyzing data, writing of the manuscript; KE, designing research studies, conducting experiments, acquiring data, analyzing data, writing the manuscript; IH, analyzing data, writing of the manuscript; BV, conducting experiments, acquiring data, analyzing data, writing the manuscript; LP, conducting experiments, acquiring data, analyzing data, writing the manuscript; SC, analyzing data, writing of the manuscript; DB, analyzing data, writing of the manuscript; EG, analyzing data, writing of the manuscript; RG, analyzing data, writing of the manuscript; SC, designing research studies, conducting experiments, acquiring data, analyzing data; HS, designing research studies, analyzing data, interpretation the results, and writing the manuscript. All authors contributed to and approved the final version of the manuscript.

