## Supplementary material for "Post-Ictal Gamma Oscillations Predict Hippocampal Structural Integrity in Mesial Temporal Lobe Epilepsy": Supplmentary figures

### Supplementary Figure 1

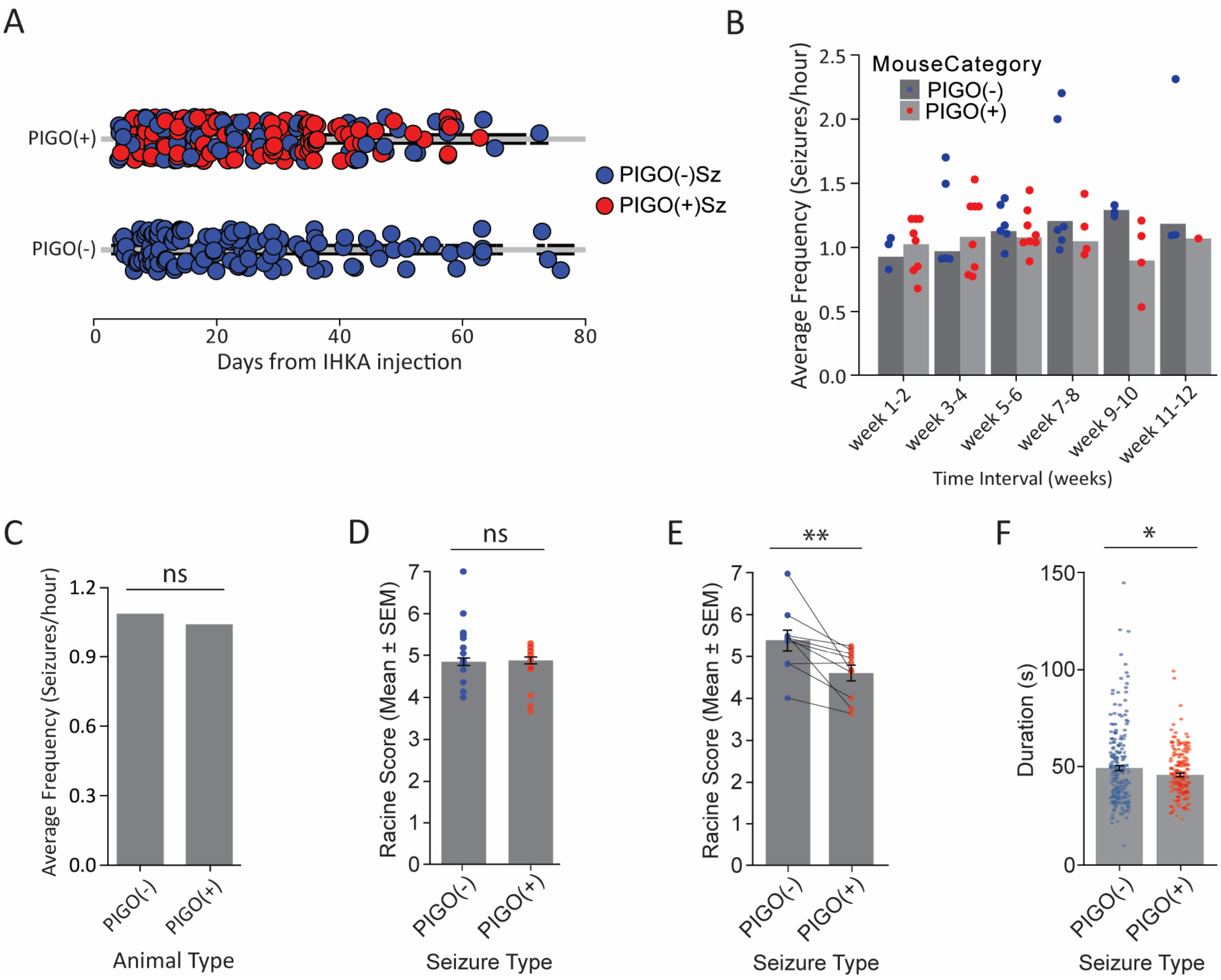


**Supplementary Figure 1. A,** Timeline of the experiment relative to the number of days after the initial KA injection, with all mice included in this study sub-grouped by PIGO(+) (Top; n=10) and PIGO(-) (Bottom; n=6) animals. **B,** Average seizure frequency of PIGO(+) and PIGO(-) animals in two-week intervals throughout the experiment. Notably, PIGO(+) animals exhibited a lower average seizure frequency after weeks 3-4 of the recording timeline compared to PIGO(-) animals. **C,** Average cumulative frequency of seizures for PIGO(+) and PIGO(-) animals (PIGO(+) = 1.04, PIGO(-) = 1.08; P=0.1, Unpaired t-test). **D,** Average cumulative Racine score of PIGO(-) and PIGO(+) seizures during the ictal period (PIGO(-) = 4.81 ± 0.09; PIGO(+) = 4.83 ± 0.08; P=0.7; Wilcoxon rank sum test). **E,** Average Racine score of during the ictal period of seizures from PIGO(+) animals only (n=10; PIGO(-)Sz = 5.39 ± 0.25, PIGO(+)Sz = 4.60 ± 0.18; P<0.01; Wilcoxon Signed Exact test). For **B,D,E,** dots are datapoints representing averages for an individual animal. **F,** Average duration between the seizure onset to offset for PIGO (-) and PIGO(+) seizures (PIGO(-) = 48.9 ± 1.62, PIGO (+) = 47.3 ± 0.94; P<0.05; Unpaired t-test). Dots represent individual seizures.

### Supplementary Figure 2

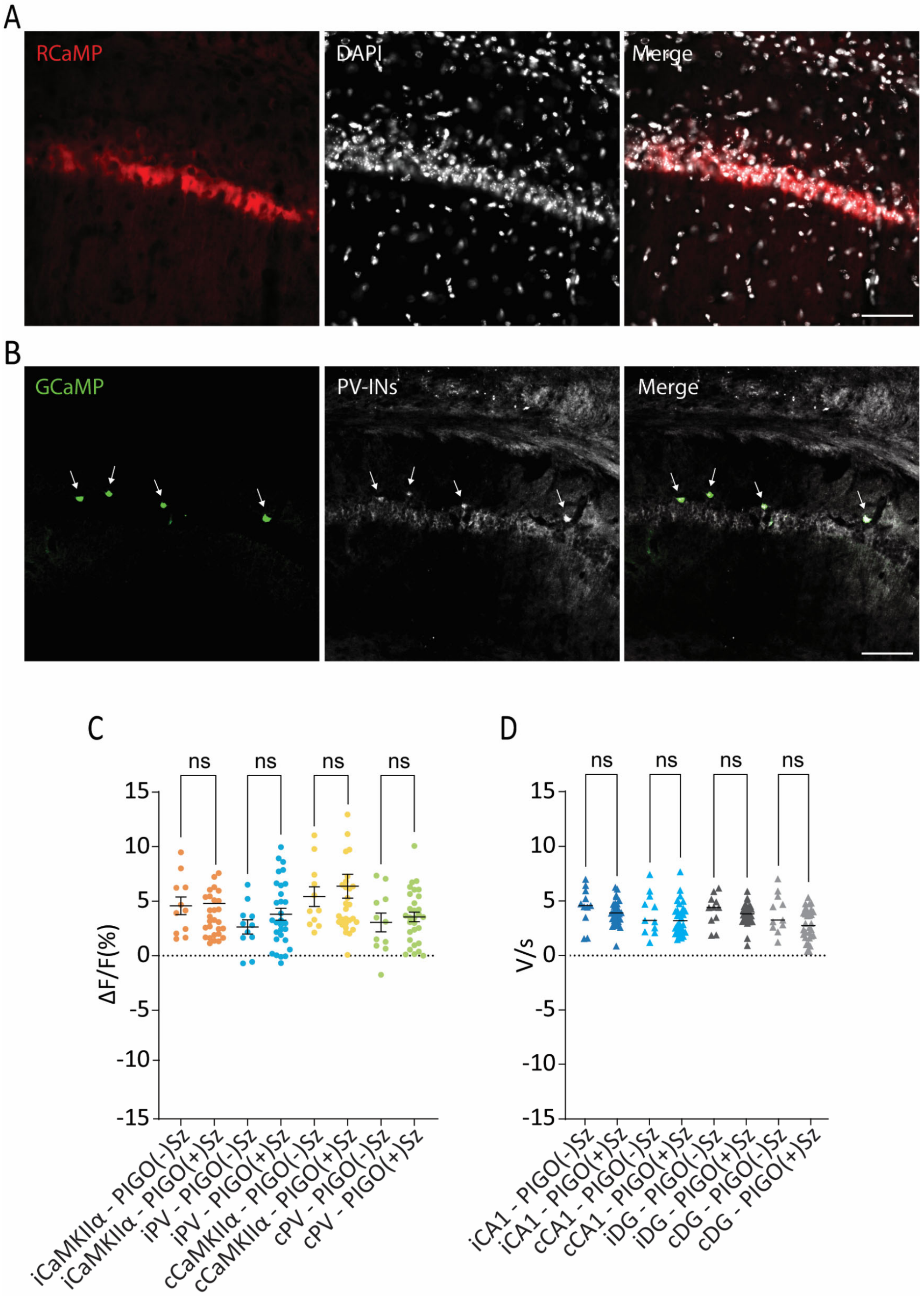


**Supplementary Figure 2. A,** Colocalization of CaMKIIα-RCaMP (red) with DAPI (white) in the CA1 layer of the antero-dorsal hippocampus. Scale Bar 50 µm. **B,** Immunofluorescence labeling for parvalbumin (PV - white) in the hippocampal CA1 layer of mice expressing the cre-dependent hSyn-GCaMP (green). Interneurons expressing hSyn-GCaMP that are parvalbumin-positive are indicated by the arrows. Scale Bars 50 µm. **C**, Analysis of $Ca^{2+}$ activity levels (ΔF/F) comparing each photometry signal (iCaMKIIα, iPV, cCaMKIIα, and cPV) obtained in the ictal period from PIGO(-)Sz and PIGO(+)Sz (iCaMKIIα-PIGO(-)Sz = 4.56 ± 0.88, iCaMKIIα-PIGO(+)Sz = 4.78 ± 0.58, P=0.89; iPV-PIGO(-)Sz = 2.52 ± 0.71, iPV-PIGO(+)Sz = 3.80 ± 0.54, P=0.22; cCaMKIIα-PIGO(-)Sz = 4.85 ± 0.77, cCaMKIIα-PIGO(+)Sz = 6.36 ± 1.09, P=0.44; cPV-PIGO(-)Sz = 2.84 ± 0.92, cPV-PIGO(+)Sz = 3.54 ± 0.44, P=0.45; Unpaired t-test). **D**, Analysis of CI levels comparing the averaged CI from each EEG signal (iCA1, cCA1, iDG, and cDG) recorded during the 60 seconds of the ictal period of PIGO(-)Sz and PIGO(+)Sz (iCA1-PIGO(-)Sz = 4.40 ± 0.57 V/s, iCA1-PIGO(+)Sz = 3.90 ± 0.20 V/s, P=0.30; cCA1-PIGO(-)Sz = 3.74 ± 0.62 V/s, cCA1-PIGO(+)Sz = 3.39 ± 0.26 V/s, P=0.56; iDG-PIGO(-)Sz = 4.07 ± 0.46 V/s, iDG-PIGO(+)Sz = 3.76 ± 0.17 V/s, P=0.45; cDG-PIGO(-)Sz = 3.78 ± 0.58 V/s, cDG-PIGO(+)Sz = 2.82 ± 0.25, P=0.09; Unpaired t-test).

### Supplementary Figure 3

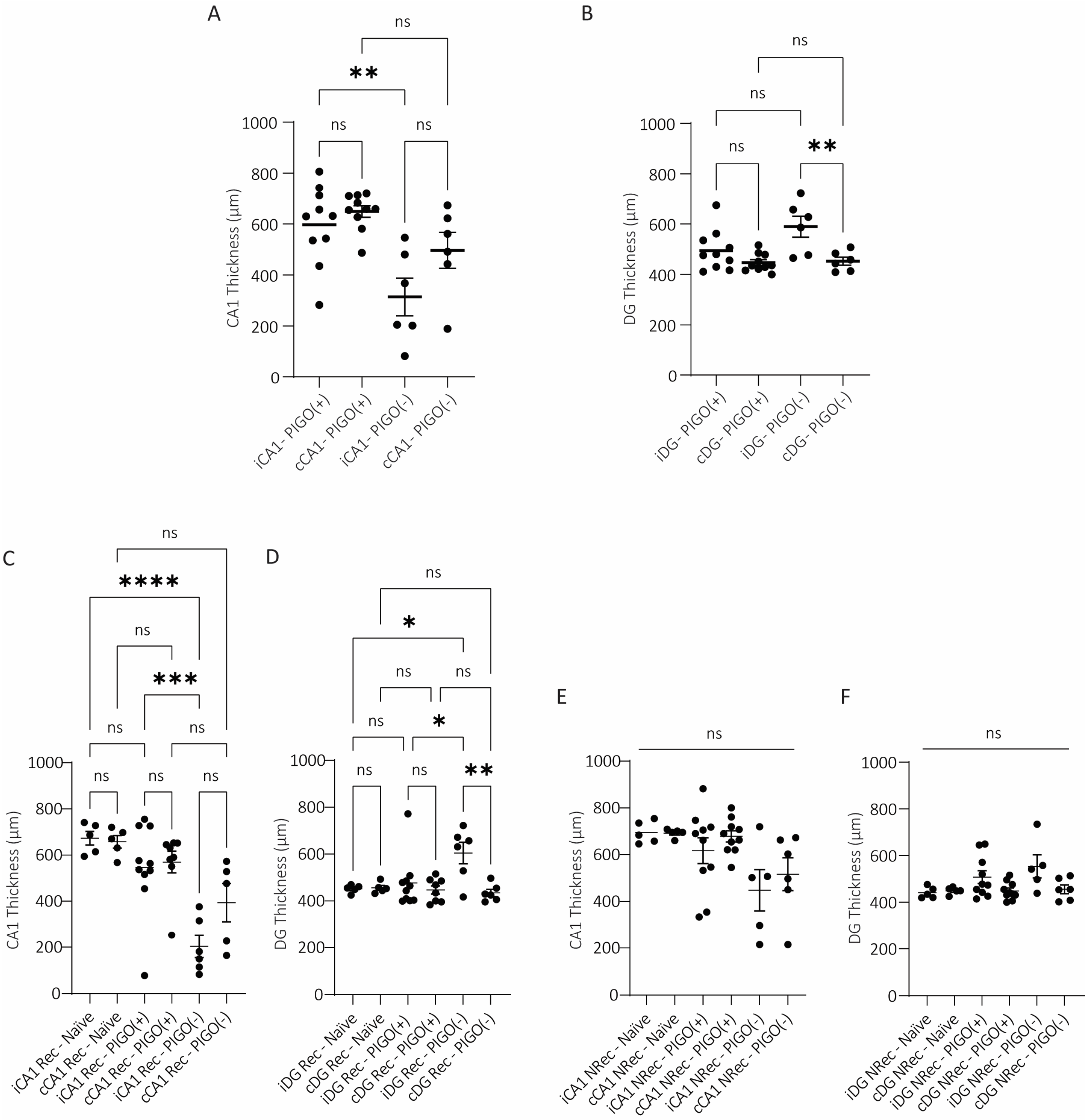

**Supplementary Figure 3. A**, Hemispheric-dependent comparison of the averaged CA1 thickness in the entire hippocampal extension of PIGO(+) and PIGO(-) animals (iCA1-PIGO(+) = 579.4 ± 48.9 µm, cCA1-PIGO(+) = 649.4 ± 22.4 µm, iCA1-PIGO(-) = 314.2 ± 73.5 µm, cCA1-PIGO(-) = 496.5 ± 70.3 µm; iCA1-PIGO(+) vs. cCA1-PIGO(+): P>0.05 ; iCA1-PIGO(-) vs. cCA1-PIGO(-): P>0.05; iCA1-PIGO(+) vs. iCA1-PIGO(-): P<0.01; cCA1-PIGO(+) vs. cCA1-PIGO(-): P>0.05; One-Way ANOVA, Tukey's multiple comparisons test as *post hoc*). **B**, As in A, but for the averaged DG thickness (iDG-PIGO(+) = 495.0 ± 25.3 µm, cDG-PIGO(+) = 447.2 ± 11.3 µm, iDG-PIGO(-) = 589.4 ± 41.5 µm, cDG-PIGO(-) = 452.9 ± 15.8 µm; iDG-PIGO(+) vs. cDG-PIGO(+): P>0.05; iDG-PIGO(-) vs. cDG-PIGO(-): P<0.01; iDG-PIGO(+) vs. iDG-PIGO(-): P>0.05; cDG-PIGO(+) vs. cDG-PIGO(-): P>0.99; One-Way ANOVA, Tukey's multiple comparisons test as *post hoc*). **C**, Hemispheric-dependent comparison of the averaged CA1 thickness in Rec areas of the hippocampus of Naïve, PIGO(+) and PIGO(-) animals (iCA1 Rec Naïve = 673.1 ± 29.4 µm, cCA1 Rec – Naïve = 657.6 ± 26.9 µm, iCA1 Rec-PIGO(+) = 546.8 ± 61.4 µm, cCA1 Rec-PIGO(+) = 569.7 ± 47.1 µm, iCA1 Rec-PIGO(-) = 203.4 ± 47.6 µm, cCA1 Rec-PIGO(-) = 394.1 ± 18.5 µm; iCA1 Rec-Naïve vs. iCA1 Rec-PIGO(+): P=0.61, iCA1 Rec-Naïve vs. iCA1 Rec-PIGO(-): P<0.0001, cCA1 Rec-Naïve vs. cCA1 Rec-PIGO(+): P=0.89, cCA1 Rec-Naïve vs. cCA1 Rec-PIGO(-): P=0.07, iCA1 Rec-PIGO(+) vs. cCA1 Rec-PIGO(+): P=0.99; iCA1 Rec-PIGO(-) vs. cCA1 Rec-PIGO(-): P=0.24; iCA1 Rec-PIGO(+) vs. iCA1 Rec-PIGO(-): P<0.01; cCA1 Rec-PIGO(+) vs. cCA1 Rec-PIGO(-): P=0.26; One-Way ANOVA, Tukey's multiple comparisons test as *post hoc*). **D**, As in C, but for the averaged DG thickness in Rec areas (iDG Rec-Naïve = 452.0 ± 7.5 µm, cDG Rec-Naïve = 456.0 ± 10.1 µm, iDG Rec-PIGO(+) = 476.7 ± 35.1 µm, cDG Rec-PIGO(+) = 446.9 ± 17.6 µm, iDG Rec-PIGO(-) = 604.6 ± 45.7 µm, cDG Rec-PIGO(-) = 434.9 ± 14.9 µm; iDG Rec-Naïve vs. iDG Rec-PIGO(+): P=0.99, iDG Rec-Naïve vs. iDG Rec-PIGO(-): P<0.05, cDG Rec-Naïve vs. cDG Rec-PIGO(+): P>0.99, cDG Rec-Naïve vs. cDG Rec-PIGO(-): P=0.99, iDG Rec-PIGO(+) vs. cDG Rec-PIGO(+): P=0.88; iDG Rec-PIGO(-) vs. cDG Rec-PIGO(-): P<0.05; iDG Rec-PIGO(+) vs. iDG Rec-PIGO(-): P<0.05; cDG Rec-PIGO(+) vs. cDG Rec-PIGO(-): P=0.99; One-Way ANOVA, Tukey's multiple comparisons test as *post hoc*). **E**, Hemispheric-dependent comparison of the averaged CA1 thickness in non-recording areas (NREC) of the hippocampus of Naïve, PIGO(+) and PIGO(-) animals (iCA1 NRec Naïve = 695.4 ± 21.3 µm, cCA1 NRec – Naïve = 692.2 ± 8.2 µm, iCA1 NRec-PIGO(+) = 617.3 ± 55.1 µm, cCA1 NRec-PIGO(+) = 678.4 ± 23.5 µm, iCA1 NRec-PIGO(-) = 448.0 ± 88.6 µm, cCA1 NRec-PIGO(-) = 516.6 ± 70.5 µm; iCA1 NRec-Naïve vs. iCA1 NRec-PIGO(+): P=0.89, iCA1 NRec-Naïve vs. iCA1 NRec-PIGO(-): P=0.06, cCA1 NRec-Naïve vs. cCA1 NRec-PIGO(+): P>0.99, cCA1 NRec-Naïve vs. cCA1 NRec-PIGO(-): P=0.28, iCA1 NRec-PIGO(+) vs. cCA1 NRec-PIGO(+): P=0.91; iCA1 NRec-PIGO(-) vs. cCA1 NRec-PIGO(-): P=0.95; iCA1 NRec-PIGO(+) vs. iCA1 NRec-PIGO(-): P=0.22; cCA1 NRec-PIGO(+) vs. cCA1 NRec-PIGO(-): P=0.21; One-Way ANOVA, Tukey's multiple comparisons test as *post hoc*). **F**, As in **E**, but for the averaged DG thickness in NRec areas (iDG NRec-Naïve = 441.3 ± 10.9 µm, cDG NRec-Naïve = 449.7 ± 6.7 µm, iDG NRec-PIGO(+) = 507.7 ± 27.4 µm, cDG NRec-PIGO(+) = 449.1 ± 11.8 µm, iDG NRec-PIGO(-) = 554.2 ± 49.6 µm, cDG NRec-PIGO(-) = 456.1 ± 18.8 µm; iDG NRec-Naïve vs. iDG NRec-PIGO(+): P=0.42, iDG NRec-Naïve vs. iDG NRec-PIGO(-): P=0.08, cDG NRec-Naïve vs. cDG NRec-PIGO(+): P>0.99, cDG NRec-Naïve vs. cDG NRec-PIGO(-): P>0.99, iDG NRec-PIGO(+) vs. cDG NRec-PIGO(+): P=0.33; iDG NRec-PIGO(-) vs. cDG NRec-PIGO(-): P=0.14; iDG NRec-PIGO(+) vs. iDG NRec-PIGO(-): P=0.77; cDG NRec-PIGO(+) vs. cDG NRec-PIGO(-): P>0.99; One-Way ANOVA, Tukey's multiple comparisons test as *post hoc*).

### Supplementary Figure 4

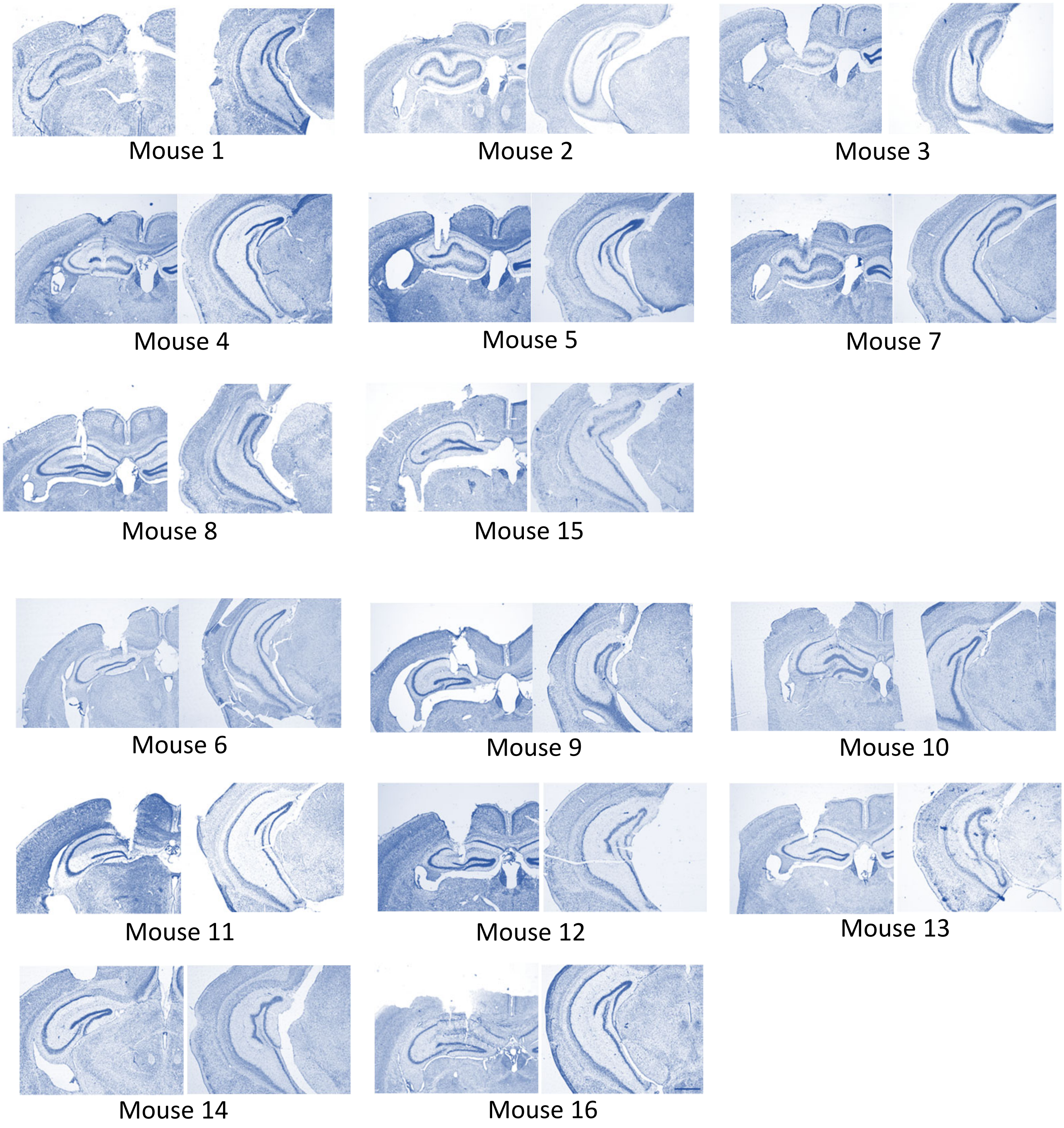


**Supplementary Figure 4.** Nissl staining photomicrographs of the hippocampus ipsilateral to the IHKA injection from all the mice included in this study. Mice 1, 2, 3, 4, 5, 7, 8 and 15 received a unilateral IHKA injection in the CA1 layer of the antero-dorsal hippocampus (adIHKA mice). Mice 6, 9, 10, 11, 12, 13, 14, and 16 received a unilateral IHKA injection in the CA1 layer of the posterior-ventral hippocampus (pvIHKA mice). For each mouse, the pictures on the left represent a brain slice containing the antero-dorsal portion of the hippocampus, while the pictures on the right show a brain slice containing the posterior-ventral portion of the hippocampus. Scale bar: 1000µm.

### Supplementary Figure 5

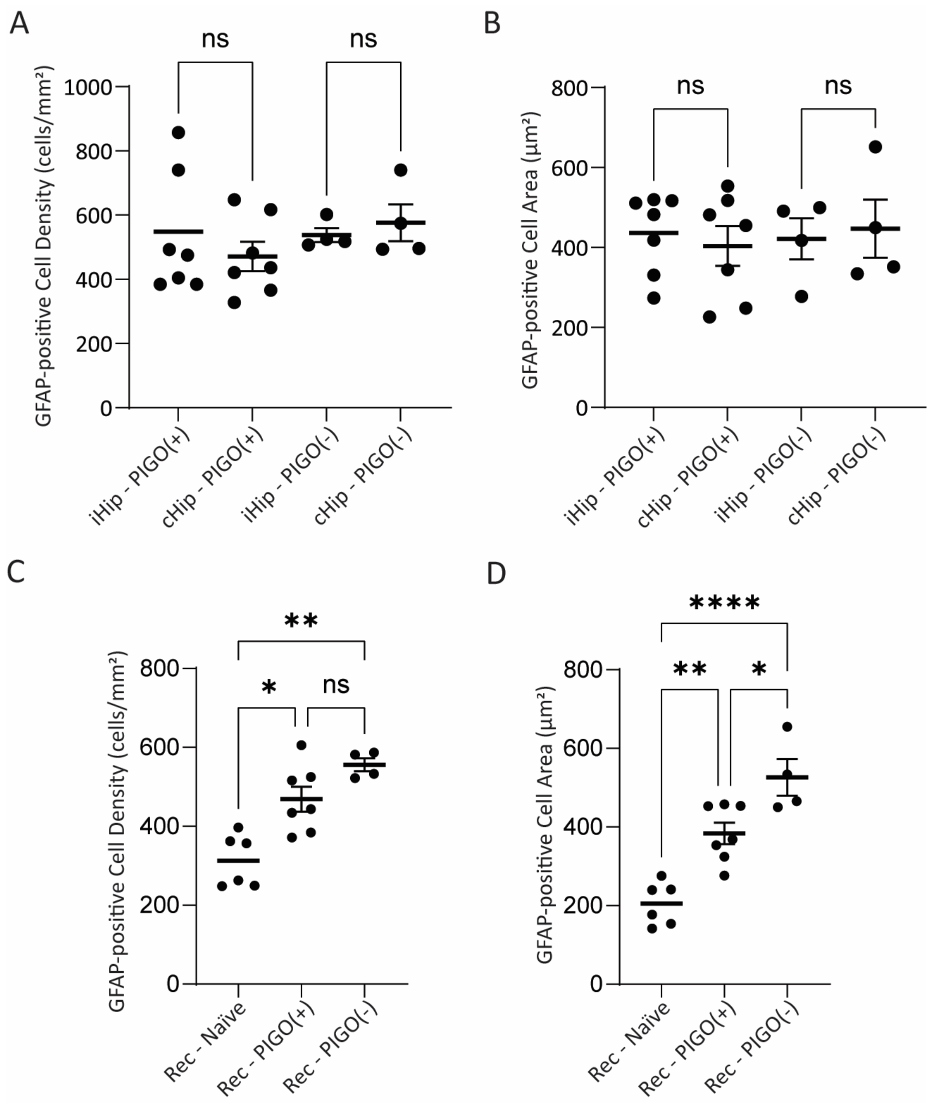


**Supplementary Figure 5. A**, Hemispheric-dependent differences in GFAP-positive cells density averages (iHip-PIGO(+) = 548.5 ± 82.78 cells/$mm^2$, cHip-PIGO(+) = 471.3 ± 45.7 cells/$mm^2$, P=0.99; iHip-PIGO(-) = 537.9 ± 21.7 cells/$mm^2$ , cHip-PIGO(-) = 576.3 ± 57.8 cells/$mm^2$, P=0.99; One-Way ANOVA, Tukey's multiple comparisons test as *post hoc*). **B**, Hemispheric-dependent differences in the area of GFAP-positive cells (iHip-PIGO(+) = of 436.2 ± 37.4 $\mu m^2$, cHip-PIGO(+) 403.8 ± 49.6 $\mu m^2$, P>0.99; iHip-PIGO(-) = 421.4 ± 51.3 $\mu m^2$, cHip-PIGO(-) = 446.8 ± 72.8 $\mu m^2$, P>0.99; One-Way ANOVA, Tukey's multiple comparisons test as *post hoc).* **C**, Averaged astrocytic density in Recording areas of the hippocampus of PIGO(+) and PIGO(-) animals in comparison to Naïve animals (Rec-Naïve = 312.8 ± 49.71 cells/$mm^2$, Rec-PIGO(+) = 448.5 ± 32.4 cells/$mm^2$, Rec-PIGO(-) = 555.8 ± 16.6 cells/$mm^2$, Rec-Naïve vs. Rec-PIGO(+): P<0.05; Rec-Naïve vs. Rec-PIGO(-), P<0.01; One-way ANOVA, Tukey's multiple comparisons test as *post hoc*). **D**, Averaged area of single astrocytes in Rec areas of the hippocampus of PIGO(+) and PIGO(-) animals in comparison to Naïve animals (Rec-Naïve 221.6 ± 22.7 $\mu m^2$, Rec-PIGO(+) = 383.2 ± 45.0 $\mu m^2$, Rec-PIGO(-) = 532.1 ± 33.2 $\mu m^2$; Rec-Naïve vs. Rec-PIGO(+): P<0.01; Rec-Naïve vs. Rec-PIGO(-), P<0.0001; One-way ANOVA, Tukey's multiple comparisons test as *post hoc*).

### Supplementary Figure 6

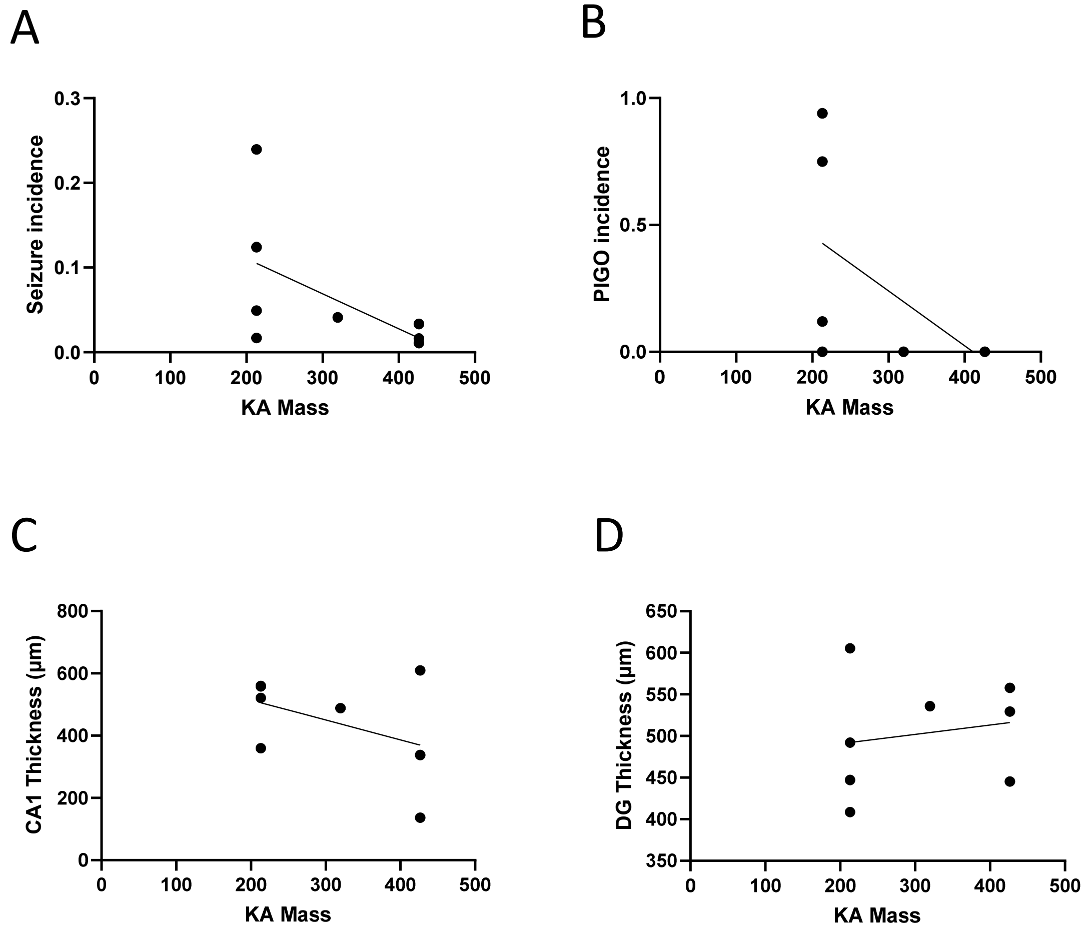


**Supplementary Figure 6. Effect of the different doses of IHKA administered to the adIHKA animals**. **A**, Seizure incidence (r=0.3073, P=0.1539); **B**, PIGO incidence (r=0.3471, P=0.1243); **C**, CA1 Thickness (r=0.1828, P=0.2907); and **D**, DG Thickness (r=0.03185, P=0.6724).

### Supplementary Table 1

| Mouse ID | Sex | Age[a] | Hours of Recording[b] | KA Injection Location | KA Mass[c] | Seizures (n) | PIGO(+)Sz[e] |
|---|---|---|---|---|---|---|---|
| M#1 | F | 102 | 244.5 | Antero-Dorsal | 2.13 | 12 | 0 |
| M#2 | M | 231 | 808.4 | Antero-Dorsal | 4.26 | 13 | 0 |
| M#3 | M | 231 | 928.4 | Antero-Dorsal | 4.26 | 10 | 0 |
| M#4 | F | 90 | 228.3 | Antero-Dorsal | 3.19 | 9 | 0 |
| M#5 | M | 89 | 209.9 | Antero-Dorsal | 4.26 | 7 | 0 |
| M#6[f] | M | 292 | 1255 | Posterior-Ventral | 2.13 | 87 | 0 |
| M#7 | F | 158 | 1547.2 | Antero-Dorsal | 2.13 | 26 | 2 (8) |
| M#8 | F | 104 | 16.7 | Antero-Dorsal | 2.13 | 4 | 3 (75) |
| M#9[f] | M | 178 | 301.9 | Posterior-Ventral | 2.13 | 6 | 4 (67) |
| M#10[f] | M | 139 | 205.2 | Posterior-Ventral | 2.13 | 16 | 12 (75) |
| M#11[f] | M | 201 | 646 | Posterior-Ventral | 2.13 | 17 | 16 (94) |
| M#12 | F | 144 | 552 | Posterior-Ventral | 2.13 | 46 | 19 (41) |
| M#13 | F | 183 | 615.4 | Posterior-Ventral | 2.13 | 33 | 24 (73) |
| M#14[f] | M | 152 | 552 | Posterior-Ventral | 2.13 | 39 | 31 (79) |
| M#15 | F | 104 | 266.1 | Antero-Dorsal | 2.13 | 33 | 31 (94) |
| M#16[f] | M | 152 | 1011.6 | Posterior-Ventral | 2.13 | 58 | 47 (81) |
| M#17[g] | M | 67 | 539.5 | Posterior-Ventral | 2.13 | 58 | 41(71) |
| M#18[g] | M | 67 | 539.5 | Posterior-Ventral | 2.13 | 41 | 25(61) |

[a] Age in days at the date of Kainic acid injection
[b] Number of total hours animal was recorded during the experiment
[c] Kainic acid mass (in ng, Mass = KA concentration x volume injected; KA = 213.23 g/mol)
[d] Number of PIGO(+) Seizures recorded
[e] Percentage (%) of PIGO(+) seizures recorded
[f] Animals included in $Ca^{+2}$ photometry analysis
[g] Animals included in direct current (DC) analysis
